# Crane fly semiochemical overrules plant control over cyanobiont in *Azolla* symbioses

**DOI:** 10.1101/2023.02.19.528669

**Authors:** Erbil Güngör, Jérôme Savary, Kelvin Adema, Laura W. Dijkhuizen, Jens Keilwagen, Axel Himmelbach, Martin Mascher, Nils Koppers, Andrea Bräutigam, Charles van Hove, Olivier Riant, Sandra Nierzwicki-Bauer, Henriette Schluepmann

## Abstract

Semiochemicals from insects that restrict plant symbiont dinitrogen fixation had not been known. Here we report on a the glycosylated triketide δ-lactone only found in *Nephrotoma cornicina* crane flies, cornicinine, that causes chlorosis in the floating-fern symbioses from the genus *Azolla*.

Cornicinine was chemically synthesized, as well as its aglycone and diastereoisomer. Only the glycosylated trans-A form was active: 500 nM cornicinine in the growth medium turned the dinitrogen-fixing cyanobacterial filaments from *Nostoc azollae* inside the host leaf cavities into akinete-like cells. Cornicinine further inhibited akinete germination in *Azolla* sporelings, precluding re-establishment of the symbiosis during sexual reproduction. It did not affect the plant *Arabidopsis thaliana* or several free-living cyanobacteria from the genera *Anabaena* or *Nostoc*. Chlorosis occurred in hosts on nitrogen with and devoid of cyanobiont. Cornicinine, therefore, targeted host mechanisms resulting in coordinate cyanobiont differentiation.

Sequence profiling of messenger RNA from isolated leaf cavities confirmed high NH_4_-assimilation and proanthocyanidin biosynthesis in this trichome-rich tissue. Leaf-cavity transcripts in ferns grown on cornicinine reflected activation of Cullin-RING ubiquitin-ligase pathways, known to mediate metabolite signaling and plant elicitation consistent with the chlorosis phenotype. Transcripts accumulating when akinetes are induced, in leaf cavities of ferns on cornicinine and in megasporocarps, were consistent with increased JA-oxidase, sulfate transport and exosome formation.

The work begins to uncover molecular mechanisms of cyanobiont differentiation in a seed-free plant symbiosis important for wetland ecology or circular crop-production today, that once caused massive CO_2_ draw-down during the Eocene geological past.

**Significance:** Coordinated differentiation of host and filamentous cyanobacteria underlies the development of ecologically important symbioses; this includes the floating ferns *Azolla* which share their wetland habitat with *Nephrotoma cornicina* craneflies containing the glycosylated triketide δ-lactone semiochemical, cornicinine. Cornicinine overrules cyanobiont differentiation thus inhibiting symbiosis N_2_-fixation and sexual reproduction; its mode of action resembles plant elicitation as suggested by transcriptional profiling of cells lining the cyanobiont cavities using a new release of the fern host genome.

## Introduction

*Azolla* is a genus of highly productive aquatic ferns in symbiosis with the N_2_-fixing filamentous cyanobacteria *Nostoc azollae* (Nostoc). Nostoc is maintained in the fern meristems and specialized leaf cavities where it fixes enough N_2_ to sustain the astonishing growth rates of the symbiosis (Brouwer et al., 2017). The ferns’ massive depositions in Arctic sediments dating from the Eocene suggest that *Azolla* ferns may have caused climate cooling (Brinkhuis et al., 2006). In the past they were deployed as a biofertilizer, today for they have great potential for the restauration of subsiding wetlands or for the circular use of mineral nutrients in sustainable agriculture toproduce high-protein feed (Schluepmann et al., 2022). Despite this and the similar importance of symbioses of filamentous cyanobacteria with mosses (Stuart et al., 2020), the mechanisms maintaining the coordinated development of cyanobiont and host are poorly understood. Here we learn from nature how a chemical from the crane fly, *Nephrotoma cornicina,* interferes with these mechanisms at nanomolar concentrations.

Morphological observations have associated secretory trichomes (ST) with symbiosis maintenance in the shoot apical meristems, upper leaf lobes and inside the sporocarps (Calvert et al., 1985; Zheng et al., 2008). At the shoot tips, the cabbage-like crop of leaves tightly conceals the important shoot-apical Nostoc colony (SANC) and large ST. The small and likely motile SANC filaments inoculate newly forming leaf initials and sporocarps, for vertical transfer of Nostoc to the next generation (Dijkhuizen et al., 2021). Inside the cavities of the upper leaf lobes, mature N_2_-fixing Nostoc filaments are typically found along with a variety of ST. Under the indusium cap of the megaspore, ST are found along with nostoc akinetes.

Molecular mechanisms maintaining plant-cyanobacteria symbioses are known to control bacterial differentiation. Nostoc from the SANC was proposed to differentiate into motile hormogonia by hormogonia-inducing factors (HIF) secreted by the shoot apical trichomes, after which the hormogonia are attracted to the trichomes inside developing leaf cavities (Cohen et al., 2002). Diacylglycerols acting as HIF on *Nostoc* species have been isolated from the symbiotic coralloid roots of *Cycas revoluta* (Hashidoko et al., 2019). Moreover, the facultative symbiont of cycads, *Nostoc punctiforme,* has been shown to be attracted to isolated *Azolla* trichomes (Cohen et al., 2002). Once Nostoc has moved inside the leaf cavity it will differentiate into filaments with heterocysts that actively fix N_2_. The leaf-cavity trichomes could be secreting hormogonia suppressing factors (HSF) to keep Nostoc in this state. Glycosylated flavonoids such as 3-deoxyanthocyanins isolated from *Azolla* and naringin have been shown to act as HSF on *N. punctiforme* (Cohen et al., 2002). Nostopeptolides secreted by *N. punctiforme* itself also act as HSF as shown by a restored phenotype when a polyketide synthase knock-out mutant, which lacks nostopeptolides and differentiates into hormogonia by default, was supplemented with nostopeptolides (Liaimer et al., 2015). At a low concentration, nostopeptolides also acted as chemoattractant. Interestingly, when in symbiosis with the plant hosts *Gunnera manicata* or *Blasia pusilla*, nostopeptolide production by *N. punctiforme* was downregulated. Plant exudates, therefore, do influence nostopeptolide production and herewith regulate the movements and state of the cyanobiont.

Sporocarp initials (SI) of *Azolla* also have trichomes that presumably attract Nostoc and thus mediate vertical transfer of Nostoc in the life cycle of the host (Perkins and Peters, 2006; Zheng et al., 2008). In SI developing into microsporocarps, the Nostoc are not entering the microsporangia and are thus eventually lost. In contrast, the megasporocarps develop a protective indusium cap under which the Nostoc accumulate and then differentiate into akinete resting stages. Akinete inducing factors may not be required for this process because filamentous cyanobacteria are known to differentiate into akinetes when resources are limited (Zheng et al., 2013). Akinetes in the megasporocarp may be limited in nutrients and light, based on their isolation from the nutritious megaspore and the light-absorbing dark indusium cap. When an *Azolla* sporeling germinates on the tiny gametophyte formed inside the megasporocarp, it pushes towards the indusium cap. When it displaces the cap and grows through the indusium chamber it develops trichomes which are thought to reestablish the SANC (Dunham and Fowler, 1987; Peters and Perkins, 2006).

The natural environment constitutes the biggest available non-random chemical library screen to research what maintains the symbiotic interaction. Insects are the largest group in the animal kingdom and they excel at recruiting microbial symbionts with special metabolic capabilities to fill an enormous range of niches (Feldhaar, 2011). Examples of processes insect symbionts help with are digestion, detoxification and antibiotic production. Insect extracts are, therefore, a promising source to discover novel chemicals (van Moll et al., 2021). A common insect found in wetlands where *A. filiculoide*s also thrives in the Netherlands is the crane fly *Nephrotoma cornicina* (de Jong et al., 2021). These crane flies possibly spend most of their life cycle as larvae in water-drenched soil feeding on detritus while the adults only appear for sexual reproduction in midsummer. Corpses of *Nephrotoma cornicina* crane flies caused chlorotic spots in *Azolla* mats (**Figure 1A**). To reveal the compound causing this phenomenon, some ten thousand adult crane flies were collected, boiled in water, and the crude extract thus obtained fractioned, then tested for bioactivity (**Figure 1B**). The bioactive fractions were pooled and a compound with maximum absorption at 254 nm, accounting for ±0.1% DW of the crane fly biomass, could be isolated. Mass spectrometry and structural analyses characterized a novel glycosylated triketide δ-lactone, named cornicinine, which was identified as the candidate molecule turning *Azolla* chlorotic (Mathieu et al., 2005). The relative activity of cornicinine stereoisomers was not clarified.

**Figure 1.**
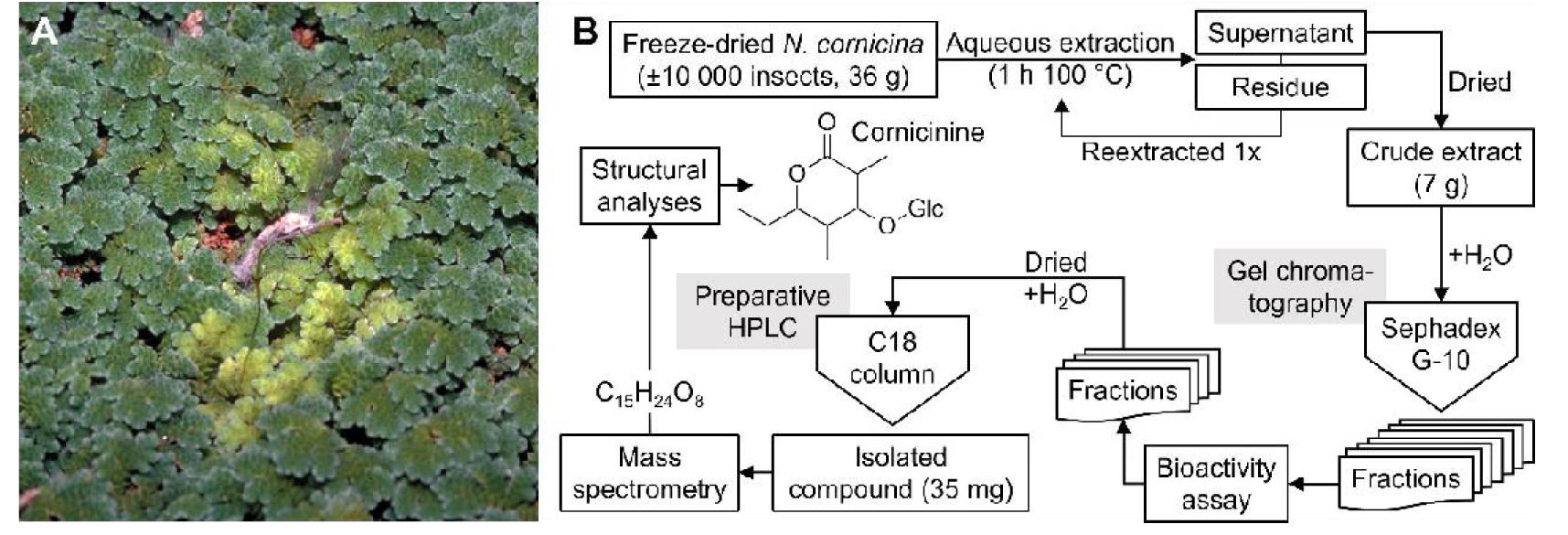
*N. cornicina* induced chlorosis in *A. filiculoides* mats and isolation procedure of cornicinine. (**A**) Typical chlorotic halo in *Azolla* mats induced by an *N. cornicina* corpse. (**B**) Overview of the procedure used to isolate, bioassay and identify cornicinine. Dry *N. cornicina* (36 g) were extracted in water, then freeze-dried yielding 7 g of dry powder. The powder was redissolved and fractioned on a Sephadex G-10 column. Fractions were assayed for bioactivity and the bioactive fractions, with a shared maximum absorption at 254 nm, were pooled. The absorption peak was used to isolate 35 mg of a pure compound by preparative HPLC. Structure elucidation by way of mass spectrometry and NMR experiments revealed a novel glycosylated triketide δ-lactone, with likely trans diastereomeric configuration, named cornicinine.

Here we examined the specific occurrence of cornicinine in insects from the genus *Nephrotoma*. We then tested chemically synthesized cornicinine stereoisomers for activity on several *Azolla* species, *Arabidopsis thaliana* and free-living *Anabaena* or *Nostoc* species. To examine the specific effect on mechanisms that control differentiation of the cyanobiont and gain first insights into the components affected by cornicinine, leaf-cavity transcripts were sequenced and compared to those in megasporocarps where *bona fide* akinetes are formed.

## Results

### *Nephrotoma cornicina* collected from around the world cause chlorosis

Corpses of the *N. cornicina* observed on the canopy of *Azolla* at the Belgium site of initial discovery were often infected with fungi (**Figure 1A, Figure S1A**). Microbes from the surrounding environment thriving on the insect biomass may therefore have been the source of the active substance. Different species of *Nephrotoma* were tested for activity including *N. aculeata, appendiculata, crocata, flavescens, flavipalpis, guestfalica, pratensis, quadrifaria, scalaris, scurra* and *submaculosa*. None of the adults from these species proved to induce chlorosis; generalist microbes on corpses from insects sharing the wetland habitat were thus not involved (**Figure S1B**). We tested *N. cornicina* individuals from Ottawa (Canada), Lucas Marsh (United Kingdom), Köyceğiz (Turkey), Segezha, Vyatka, Krasnoyarsk, Irkutsk and Sakhalin (Russia), and Kyushu (Japan), all of which displayed activity (**Figure S1C**). We thus concluded that the compound is not synthesized by microbes recruited from the environment but is systematically associated with *N. cornicina*.

### Only the trans-A diastereoisomer of cornicinine turns all tested Azolla species chlorotic; its aglycone does not

To verify the identity and activity of cornicinine purified from *N. cornicina*, two stereoisomers were synthesized chemically: the trans-A (with the R,R-lactone) and trans-B (with the S,S-lactone) (**Figure 2A**). The aglycone lactones were synthesized from the commercially available propionyl oxazolidinone stereoisomeric precursors, then glucose was added with acetylated hydroxyl groups to direct condensation reactions, and resulting acetylated intermediates were deacetylated (**Figure 2A, Figure S2)**. Key to the synthesis of the aglycone lactone, which was achieved in three steps, were the conditions to generate the Evans anti-adduct and its subsequent high-yield (71%) intramolecular lactonization in an excess of KHMDS at −78°C (**Figure S2A**). Overall, the yields for aglycone lactone synthesis were similar for the stereoisomers: 37% and 34%, respectively, for cornicinine and its diastereoisomer (**Figure S2B-C**). The O-glycosylation (68% yield) and de-acetylation (81% yield) had higher combined yields.

**Figure 2.**
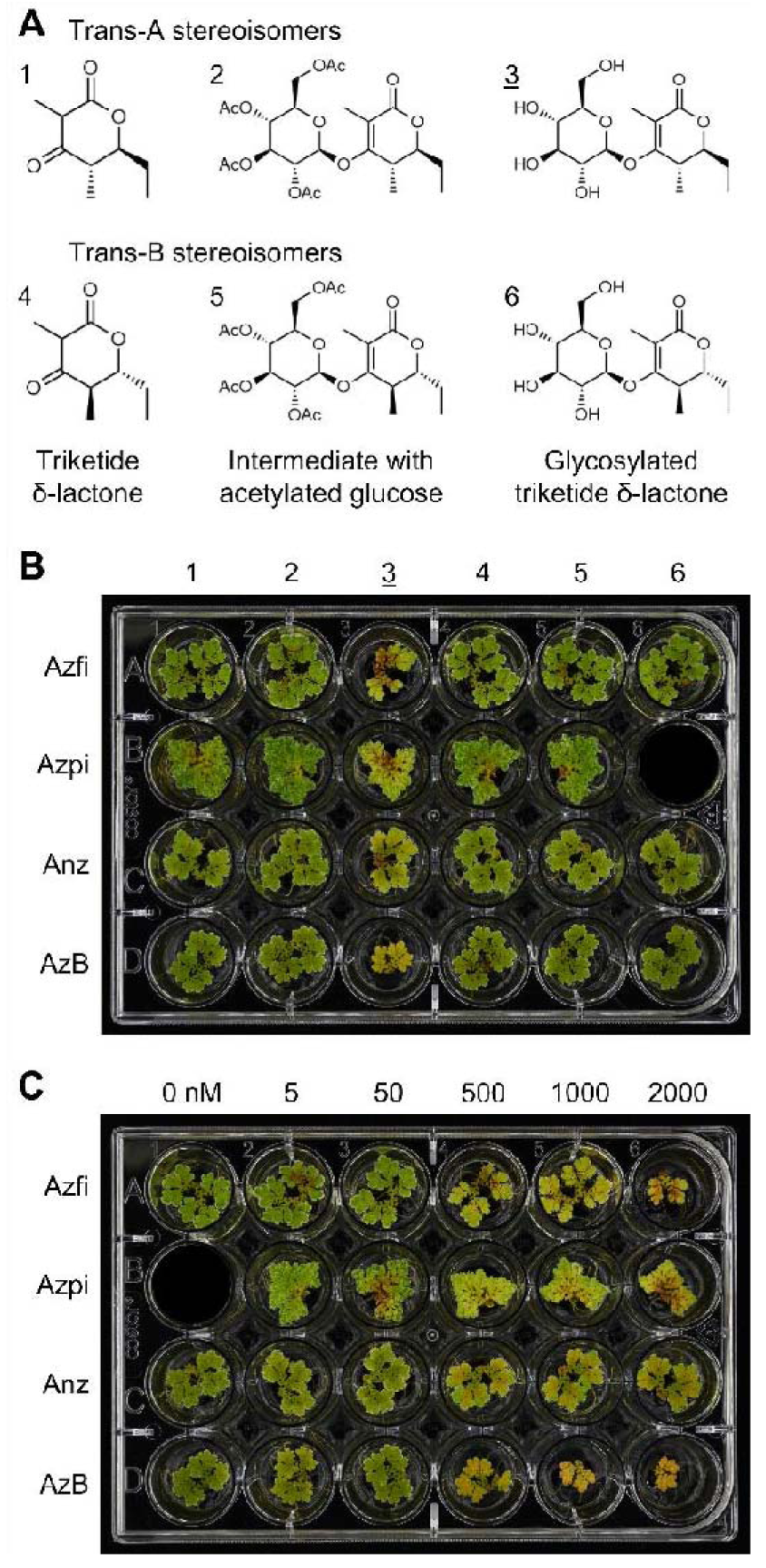
Effect of synthetic trans-stereoisomers of cornicinine, their aglycones and acetylated precursors on four *Azolla* species. (**A**) Overview of the compounds chemically synthesized in trans-A and trans-B configuration, respectively: compound 1 and 4 are the aglycones, compound 2 and 5 are the synthesis intermediates with an acetylated glucose and compound 3 and 6 are cornicinine and its diastereoisomer. (**B**) fern fronds from *Azolla* species after 25 days on 500 nM of the compounds from (**A**). (**C**) fern fronds from *Azolla* species after 25 days on a concentration gradient from 0-2000 nM of the bioactive trans-A diastereoisomer cornicinine, compound <u>3</u> in (**B**). Azfi: *A. filiculoides*; Azpi: *A. pinnata*; Anz: *Azolla* species from Anzali (Iran); AzB: *Azolla* species from Bordeaux (France).

For both stereoisomeric forms, the aglycone, acetylated synthesis intermediate and cornicinine were then supplemented, at a concentration of 500 nM, to growth medium with shoot tips of four different *Azolla* species representing both sections of the *Azolla* genus: *Azolla* and *Rhizosperma*. After 25 days, the ferns supplemented with trans-A cornicinine were chlorotic but not those with trans-B cornicinine (**Figure 2B**). The first signs of yellowing and growth retardation were already visible after 6 days and gradually increased over time (**Figure S3**). The aglycone did not cause chlorosis, proving that glycosylation is essential for the bioactivity of trans-A cornicinine (from now on referred to as cornicinine). The acetylated compounds also had no activity.

When testing cornicinine concentrations ranging from 5 nM to 2000 nM, 500 nM cornicinine generally sufficed to cause chlorosis in all species tested (**Figure 2C**). *A. filiculoides* and *A. pinnata* turned yellow and stopped growing gradually over time when on 500-2000 nM cornicinine (**Figure 2C, Figure S4**). *Azolla* sp. Bordeaux was the most sensitive with severe growth retardation at 1000 nM cornicinine and above. *Azolla* sp. Anzali was affected the least with similar growth rate and phenotype at 500-2000 nM cornicinine. Both, the species from Bordeaux and Anzali started turning red consistent with 3-deoxyanthocyanin accumulation after 17 days (**Figure S4**). The chlorosis in combination with growth retardation made us wonder what is happening to the cyanobacterial symbiont.

### Cornicinine induces the coordinate differentiation of N. azollae filaments from the leaf cavities into akinete-like cells within six days

We visualized Nostoc by crushing shoot tips between two glass slides for microscopy after 12 days growth with 500 nM synthetic compound. Cornicinine-treated *A. filiculoides* fern fronds did not have the heterocyst-rich Nostoc filaments characteristic of leaf cavities (**Figure 3A**). Instead, single, larger cells with cyanophycin granules accumulated that resembled the akinetes found under the indusium cap of the megasporocarp (**Figure S5**). The cornicinine-induced akinetes had a more elongated shape than those typical of the indusium and we therefore called them akinete-like cells (ALC). The shapes of the ALC of the four tested *Azolla* species looked surprisingly different (**Figure 3B**). The ALC of *A. filiculoides* mostly contained two to five cyanophycin granules while the ALC of *A. pinnata* were smaller and did not seem to contain any granules. The ALC of the *Azolla* species from Anzali and Bordeaux were larger, sometimes rhombus shaped, and contained five to ten granules.

**Figure 3.**
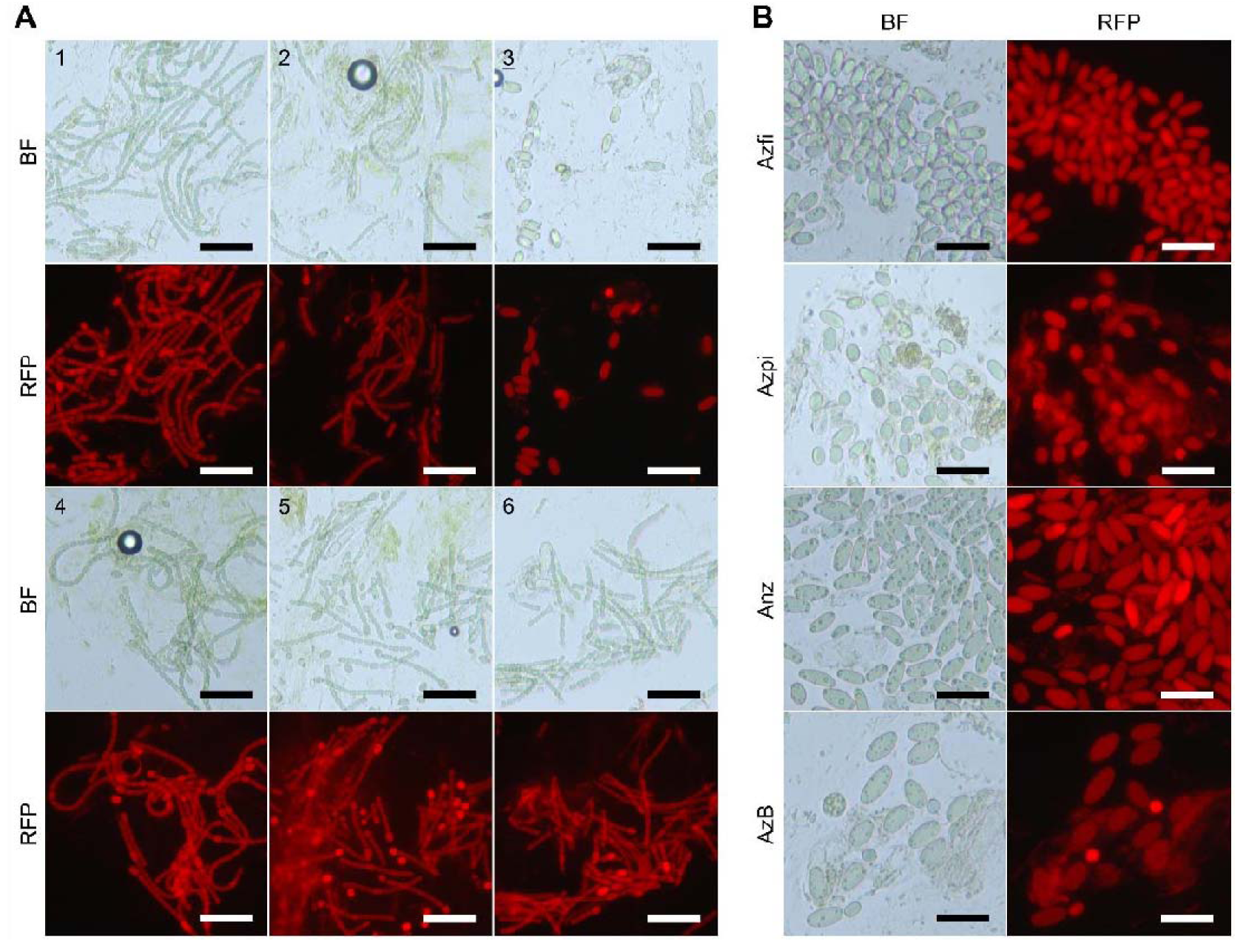
Effect of synthetic trans-stereoisomers of cornicinine, their aglycones and acetylated precursors on *N. azollae* from four *Azolla* species. (A) *N. azollae* inside *A. filiculoides* after 12 days on 500 nM of the six compounds from Figure 2A, scale bars correspond to 50 μm. (B) Morphology of the akinete-like cells from four different *Azolla* species induced by 500 nM cornicinine after 12 days, scale bars correspond to 30 μm. The different *Azolla* species were as in Figure 2. BF: bright-field; RFP: fluorescence under the settings for red fluorescent protein (RFP).

The trans-B stereoisomer and aglycone of cornicinine neither induced ALC, chlorosis nor growth retardation. Development of ALC thus had to be a result of cornicinine. We followed the development of ALC over time and with a range of concentrations. After 6 days on more than 1000 nM cornicinine, all Nostoc from *A. filiculoides* fern fronds were differentiated into ALC, while on 500 nM cornicinine isolated filaments were still present albeit with somewhat bloated vegetative cells (**Figure S6**). These bloated filaments eventually completely disassociated into ALC between day eight and 12. Fern fronds treated with ≤50 nM cornicinine still contained filaments even after 21 days indicating that a threshold concentration is required before ALC are formed (**Figure S6**).

The first signs of yellowing and growth retardation after six days on 500 nM cornicinine coincided with the first signs of ALC-induction. A slight delay persisted, however, between the complete disappearance of filaments to the extent that only isolated ALC remain (day 12) and chlorosis with growth retardation (day 17-21) (**Figure S4, Figure S6)**. ALC unlikely fix N_2_, as this process is usually attributed to heterocysts fueled by the metabolism of vegetative cells in the intact Nostoc filament. The ferns may have temporarily relied on internally stored nitrogen in the time between complete ALC-induction and yellowing. Consequently, chlorosis may be a result of nitrogen starvation.

### Cornicinine-induced chlorosis is not alleviated by nitrate supplied in the medium

*A. pinnata*, *A. filiculoide*s and a strain of *A. filiculoides* devoid of Nostoc (Brouwer et al., 2017) were grown with(out) 500 nM cornicinine and 1 mM KNO_3_. After 17 days, *A. pinnata* growth inhibition by cornicinine was suppressed by nitrate, but the chlorosis caused by cornicinine remained (**Figure 4A, Figure S7**). Nitrate neither suppressed the growth inhibition nor the chlorosis of *A. filiculoides* on cornicinine regardless of the presence of the cyanobacteria. We conclude that chlorosis was caused by something else than solely the nitrogen deficiency resulting from complete differentiation of the Nostoc into ALC. We next wondered whether cornicinine affects plants or free-living cyanobacteria.

**Figure 4.**
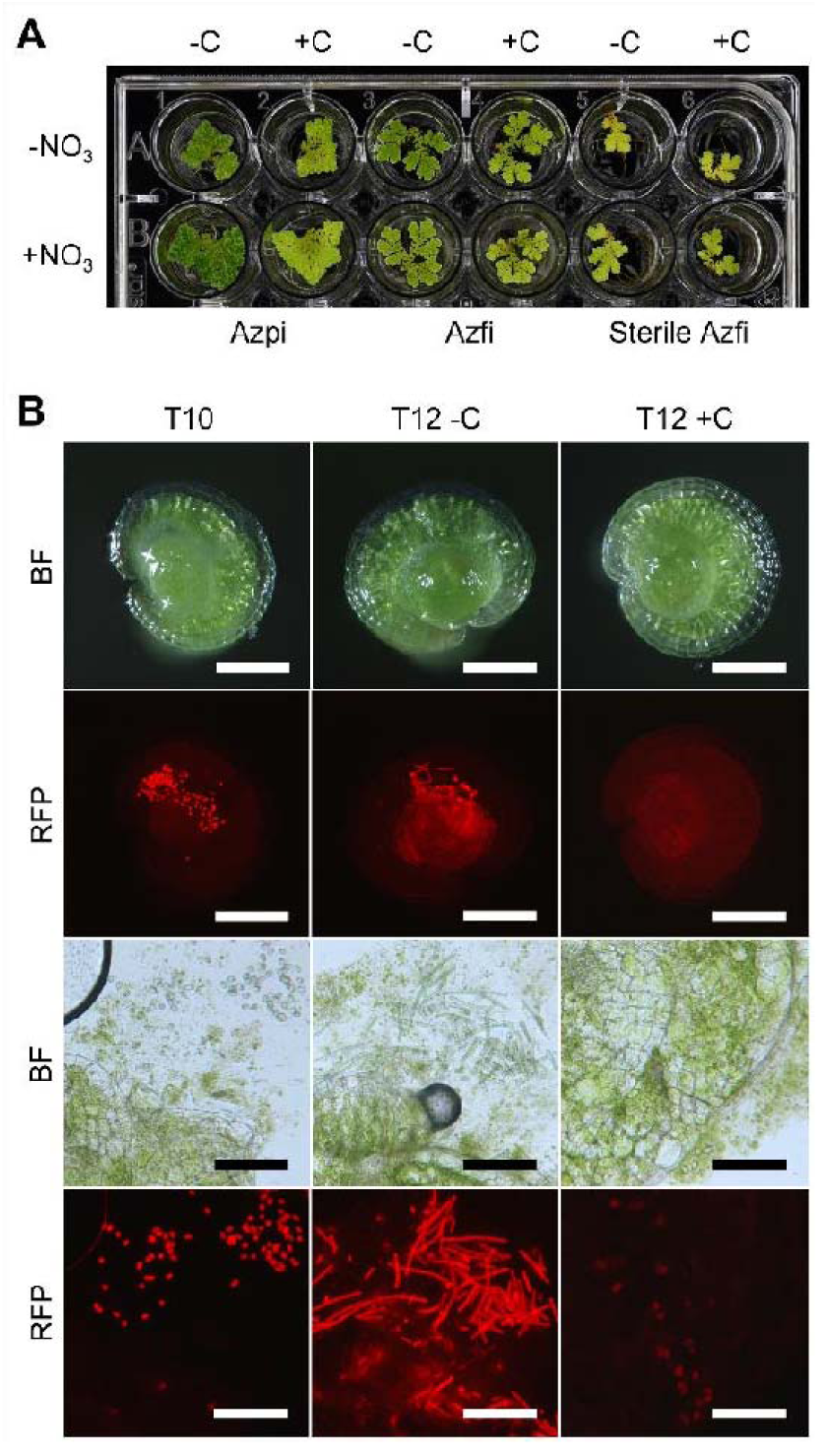
Effect of cornicinine when sporophytes grow on nitrate supplemented medium and during the reestablishment of the symbiosis when sporelings germinate. (A) *A. pinnata*, *A. filiculoides* and *A. filiculoides* devoid Nostoc after 17 days without (-C) and with 500 nM cornicinine (+C) and 1 mM KNO_3_ (-/+NO_3_). (B) Top: from left to right, *A. filiculoides* sporeling 10 days after germination, and sporelings 12 days after germination with(out) 500 nM cornicinine (-/+C). Bottom: the same sporelings crushed to expose *N. azollae*. Images are representative for 15 individual sporelings imaged per condition. Scale bars on top and bottom, respectively, correspond to 200 μm and 100 μm. The different *Azolla* species were as in Figure 2. BF: bright-field; RFP: fluorescence under RFP settings.

### Cornicinine did not affect the growth and differentiation of Arabidopsis seedlings or free-living filamentous cyanobacteria

Given the overall similarity of cornicinine to sugar disaccharides, we tested the germination and growth of Arabidopsis seedlings on medium containing 500 nM cornicinine with(out) 100 mM sucrose, or 100 mM sorbitol osmoticum control. Whilst the seedlings responded to the sucrose and osmoticum, the 500 nM cornicinine did not have visible effects on germination time, and root or shoot growth-rates and -habit under any of the conditions tested (data shown for seedlings without sugars in **Figure S8A**). Similarly, 500 nM cornicinine did not alter the growth rates of *Anabaena* PCC 7210 (**Figure S8B**); it also did not induce the differentiation into akinetes in strains of an *Anabaena sp.*, *Nostoc spugimena* and *N. punctiforme* when tested in the nitrogen-free BG-11_0_ medium (**Figure S8C**). Therefore, cornicinine interferes with mechanisms specific for the symbiosis.

### Cornicinine inhibits the germination of akinetes from the megasporocarp during sporeling germination

To test whether cornicinine would affect the dedifferentiation of *bona fide* akinetes and sporeling germination, clumps of *A. filiculoides* spores were germinated in demineralized water with(out) 500 nM cornicinine. After 10 days, the first green sporelings popped up to the water surface and were all inoculated with akinetes, suggesting that cornicinine did not interfere with germination of the fern host (**Figure 4B**). Some akinetes were enclosed by the first emerging leaf, but the majority were just attached to the outer surface of the sporeling (**Figure 4B**, T10). The latter does not need motile *Nostoc* cells, it could have resulted from the sporeling growing through the indusium and engulfing the akinetes from under the indusium cap with its cup-shaped first leaf. After 12 days, the akinetes captured by sporelings not exposed to cornicinine proliferated into filaments while those from sporelings growing with cornicinine remained akinetes (**Figure 4B**, T12 -C vs. T12 +C). After 21 days, the sporelings without cornicinine had already reached the four-leaf stage while those with cornicinine only reached the two-leaf stage (**Figure S9**). Cornicinine-treated sporelings did not exhibit the typical fluorescence under the RFP-settings compared to untreated sporelings: when crushed between two glass slides, however, akinetes were still found (**Figure S9**). The sporelings on cornicinine, therefore, failed to reestablish the symbiosis. Cornicinine inhibition of the germination of *bona fide* akinetes from Nostoc meant that it interferes with processes controlling the differentiation of the symbiont. The results further indicated that the ALC are akinetes. We next researched the physiological response to cornicinine by sequencing fern transcripts from the cells lining the leaf cavities.

### Profiles of polyA-enriched RNA from leaf-cavity preparations are consistent with expected metabolic activities in cells lining the leaf cavities

Since cornicinine suppressed the germination of Nostoc akinetes, we reasoned that cornicinine interference was a lasting state, lasting well over 6 days. We thus profiled RNA in ferns treated with(out) 500 nM cornicinine for six days, before the Nostoc akinetes were homogenously induced (**Figure 5**). Preparations from leaf cavities were highly enriched in trichomes of the leaf cavities presumed to mediate fern-cyanobiont interactions (**Figure 5B**). Despite poly-A enrichment a large proportion of read pairs sequenced from the leaf-cavity samples consisted of multi-mappers aligning to the Nostoc genome, mostly stemming from rRNA. As a result, the read pairs that mapped in a unique location to the fern genome varied from 1.3-7.5 million PE reads for the leaf-cavity samples (**Figure 5D**). In contrast, typically 75% of the PE reads from the sporophyte mapped to the fern genome in a unique location. Nevertheless, known functions of the 30 transcripts accumulating most highly and specifically in leaf-cavity samples were consistent with activities expected from cells lining the leaf cavities compared to sporophytes (**Figure 5E**, sp vs lp): reduced photosynthesis, increased secondary metabolism and transport.

**Figure 5.**
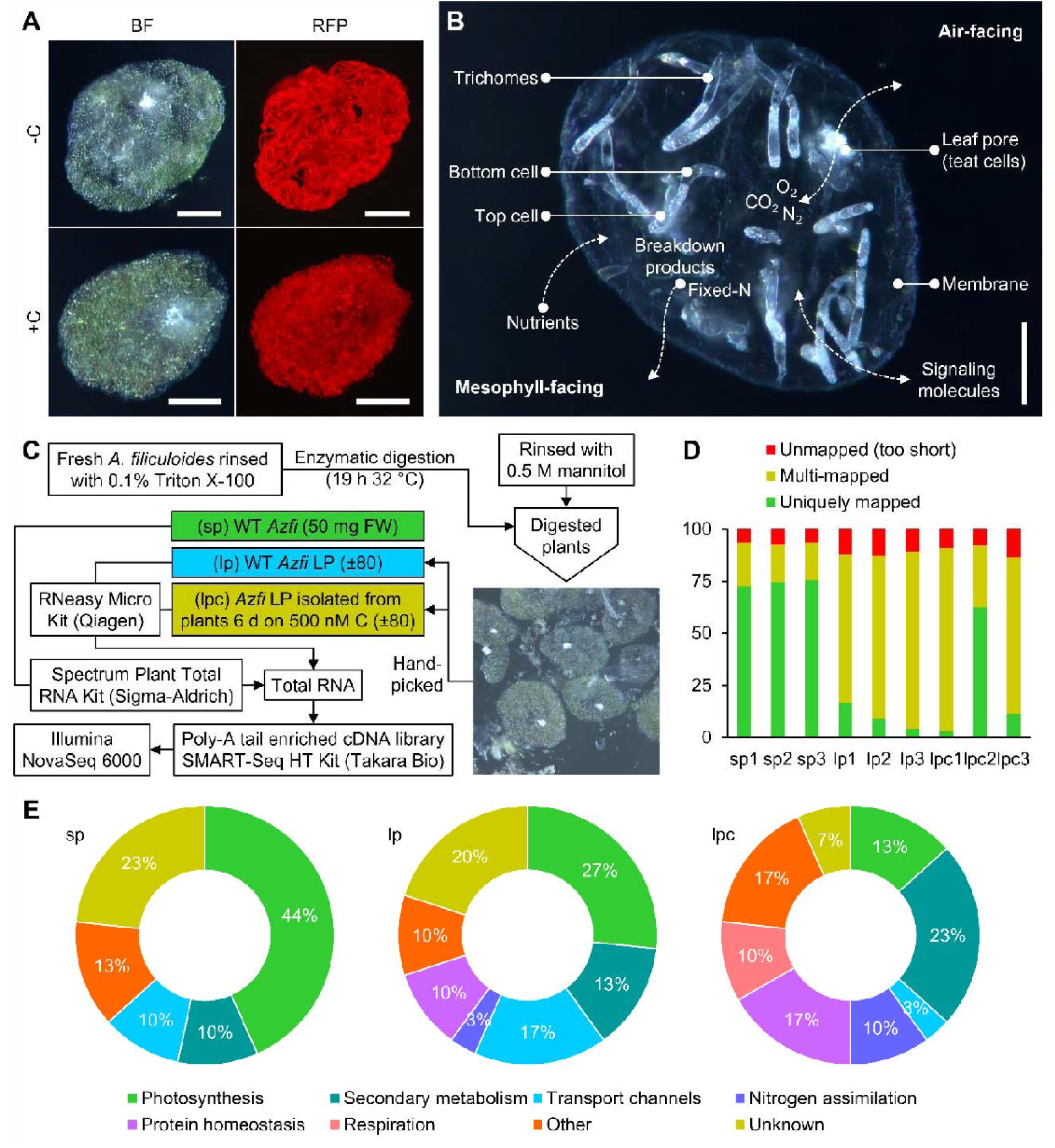
Transcription profiles of cells lining the leaf cavity in *A. filiculoides*. (**A**) leaf cavities prepared from sporophytes grown without (-C) and with 500 nM cornicinine (+C) for 10 days. BF: bright-field; RFP: fluorescence under RFP settings. Scale bars correspond to 50 μm. (**B**) Morphology of an empty leaf cavity prepared from *A. filiculoides* devoid Nostoc. The leaf pore with characteristic teat cells is air-facing while the trichomes emerge from the mesophyll-facing side. Scale bar corresponds to 100 μm. (**C**) mRNA profiling of sporophyte (sp), leaf cavities (Lp) and leaf cavities isolated from cornicinine-treated ferns (lpc). Sporophytes were treated 6 days with(out) 500 nM cornicinine; the leaf cavities were released enzymatically, then manually collected in three independent replicates per condition. All samples were collected snap frozen 2-3 h into the light period. Total RNA was extracted, DNase treated, enriched for poly-A tail before library preparation and sequencing. (**D**) Proportion of paired-end reads aligning to the concatenated genomes of *A. filiculoides*, its chloroplast, and *N. azollae* using default settings of STAR aligner. (**E**) Functional categories of the 30 genes with highest transcript accumulation per sample type.

Transcripts accumulating very highly in the leaf-cavity profiles encoded enzymes critical for N-assimilation. A cytosolic glutamine synthetase, specifically expressed in the leaf cavities, had the twelfth most read counts in the leaf-cavity profile (**Figure 6A**, GS1 Afi_v2_s3215G000080.1). An asparagine synthetase (**Figure 6A**, ASN Afi_v2_s35G001910.1) and an NADH-dependent glutamate synthase (**Figure 6A**, GOGAT Afi_v2_s35G000930.1). In contrast, transcripts of nitrate reduction were little expressed, consistent with reports identifying ammonium as the likely metabolite secreted by Nostoc (Ray et al., 1978). The accumulation of the amino acid transporter LHT (**Figure 6A**, LHT Afi_v2_s23G003000.2) could reflect amino-acid export from the cells lining the leaf cavity. Moreover, leaf-pocket profiles had very high read counts for transcripts from key enzymes of the proanthocyanidin biosynthesis pathway known to be very active in trichomes lining the leaf pocket (Güngör et al., 2021; Pereira and Carrapiço, 2007; Tran et al., 2020): the leucoanthocyanidin reductase which had the second highest read counts in the leaf-pocket profiles (**Figure 6B**, LAR Afi_v2_s74G000210.2), and two 2-oxoglutarate-dependent dioxygenases resembling anthocyanidin synthase (**Figure 6B**, Afi_v2_s16G002830.1, 2OGD-s16 and Afi_v2_s44G002700.1, 2OGD-s44) that were specifically expressed in the leaf cavities. The extraordinary numbers of reads from the SLAC (**Figure 6B**, SLAC Afi_v2_s20G000170.2) and the α-carbonic anhydrase (**Figure 6B**, α-CA Afi_v2_s189G000110.2) are reminiscent of guard cell metabolism. Given the absence of AMT transcripts, and the low PIP2 transcripts in the leaf-cavity profiles compared to whole fern (**Figure 6C, Figure S10**), NH_4_^+^/NH_3_ import may rely on the pH of the leaf cavity (**Figure 6C**). Alternatively, an as yet uncharacterized transporter of cations imports NH_4_^+^, or an altogether alternative mechanism may be involved given the strikingly high and leaf pocket specific accumulation of the NRT1/PTR (Afi_v2_s47G001780.1) transporter and NRT3 (Afi_v2_s12G001250.1) generally associated with nitrate transport and its regulation for a system where NO_3_^-^ is thought to be absent (**Figure 6A, C**).

**Figure 6.**
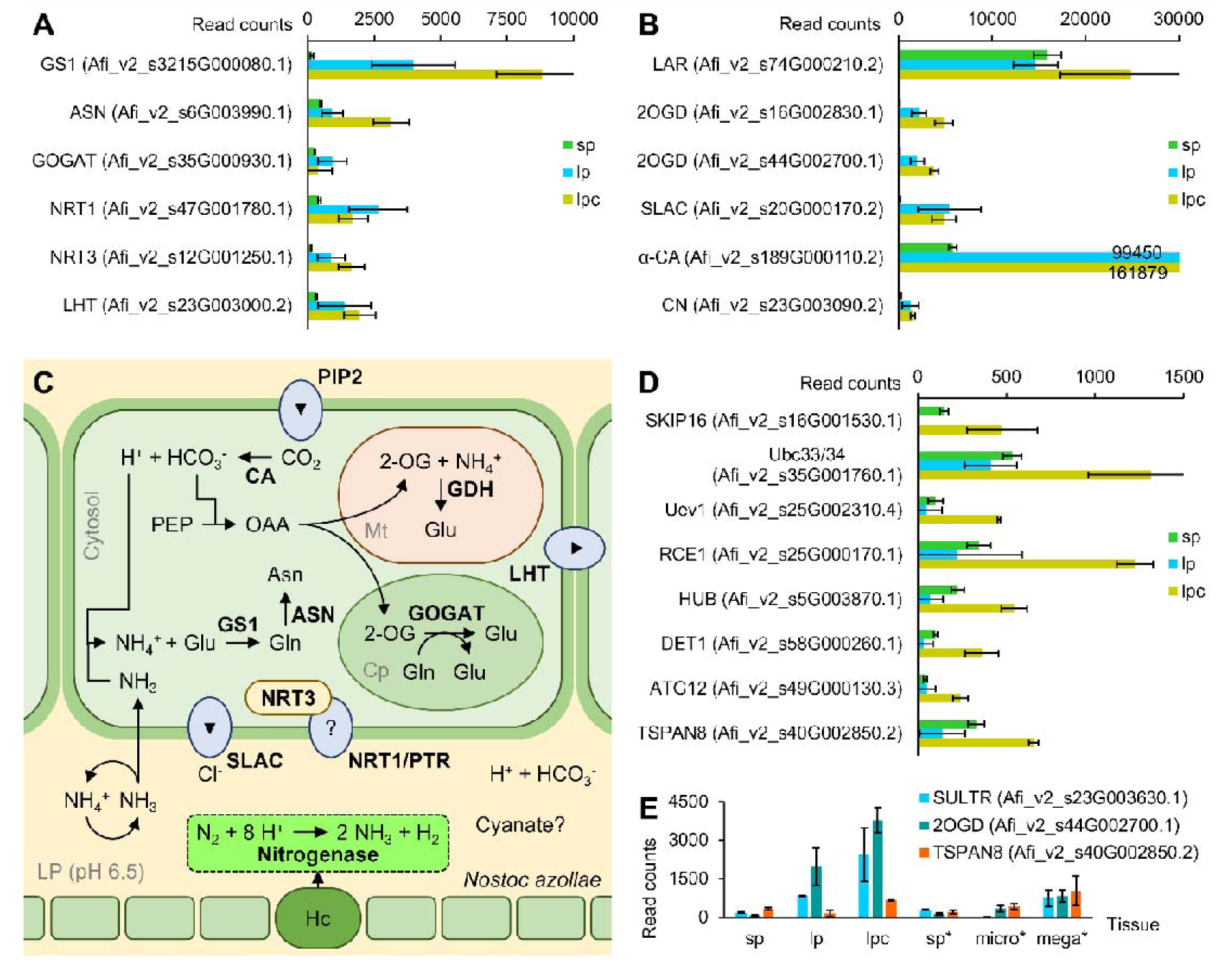
Abundant leaf-cavity transcripts related to nitrogen uptake, secondary metabolism, and responsive to cornicinine. (**A**) Ammonium assimilation and transport of nitrogenous products. (**B**) Secondary metabolism and CO_2_ solvation. (**C**) Proposed pathway for NH_4_^+^/NH_3_ assimilation in fern cells lining the leaf cavity. (**D**) Cornicinine responsive transcripts encoding F-box and ubiquitin ligase components or vesicle trafficking. α-CA, α-carbonic anhydrase; GS1, glutamine synthetase; ASN, aspartate aminotransferase; GDH, glutamate dioxygenase; GOGAT, glutamine oxoglutarate aminotransferase; SLAC, slow anion channel; LHT, neutral amino acid or ACC transporter; PIP2, plasma membrane intrinsic protein; NRT1/PTR transporters for NO_3_^-^, peptide or other solute. (**E**) Leaf-cavity specific transcripts responsive to cornicinine and upregulated in megasporocarps. *The samples were from a separate experiment profiling sporophytes (sp*), microsporocarps (micro*) and megasporocarps (mega*). Samples were collected as triplicate biological replicates 2 h into the 16 h light period. Standard deviations are shown for n=3, except for lpc where n=2.

Having established a sense of trust in the noisy signal from the leaf-pocket profiles, we proceeded with comparing the leaf-pocket profiles obtained from ferns grown with(out) cornicinine.

### Fern transcripts accumulating when akinetes are induced

Transcripts accumulating robustly in leaf cavities from ferns grown with compared to without cornicinine were few; this was in part due to the dispersion in the data and the lower sensitivity of the leaf-pocket RNA-sequencing assay (**Figure 5D**). The robust accumulation of transcripts encoding several components of the Cullin-RING ubiquitin ligase (CRUL) complexes in leaf cavities from ferns grown with cornicinine was, therefore, striking (**Figure 6D**). These components included F-box protein SKIP16-like (Afi_v2_s16G001530.1), and components of E2 and E3 ligases including the ATG12-like protein (Afi_v2_s49G000130.3) known to be involved in autophagy.

To test whether accumulation of the transcripts in leaf cavities of cornicinine grown ferns was associated with the formation of *bona fide* akinetes, we further compared their accumulation in RNA profiles from megasporocarps, compared to sporophytes (**File S1)**. This identified the sulfate transporter (Afi_v2_s23G003630.1), 2OGD-s44 (Afi_v2_s44G002700.1) and tetraspanin 8 (Afi_v2_s40G002850.2) as loci with high expression associated with *Azolla* tissues lining Nostoc akinetes (**Figure 6E**). The DOXC-class enzyme 2OGD-s44 was of particular interest since such enzymes may catalyze reactions in flavonoid biosynthesis. Only two cornicinine-induced DOXC (Afi_v2_s16G002830.1 and Afi_v2_s44G002700.1) were highly and specifically expressed in the leaf pocket, with only 2OGD-s44 also significantly expressed in megasporocarps (**Figure 7A**).

**Figure 7.**
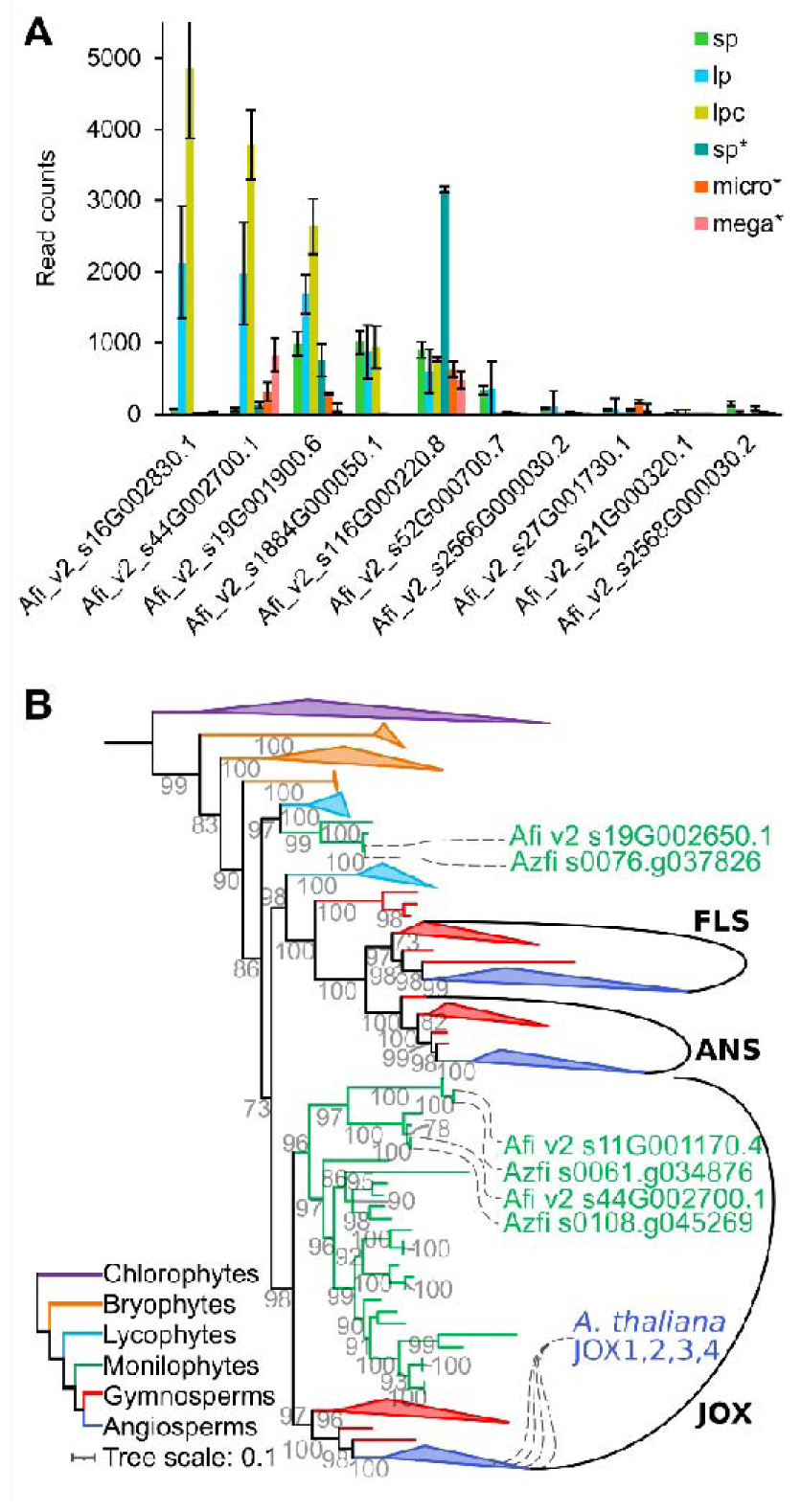
Expression and phylogeny of the DOXC enzymes from *Azolla*. (**A**) Ten most *Azolla* DOXC expressed in the leaf cavities. Samples were as in Figure 6E. (**B**) Phylogeny of 2OGD genes encoding FLS, ANS and JOX across land plant lineages. An initial phylogeny (**Figure S11**) was computed to place *A. filiculoides* genes in the broad 2OGD phylogeny. From this broad phylogeny, FLS, ANS, JOX and outgroup sequences were selected to compute a more accurate tree. Sequences were aligned with MAFFT-linsi (Katoh et al., 2019), and then trimmed using trimAL (Capella-Gutierrez et al., 2009). The phylogeny was computed with IQ-tree (Nguyen et al., 2015) with 200 non-parametric bootstraps and transfer-bootstrap values were calculated with booster (Lemoine et al., 2018).

Our phylogenomic analyses, however ascertained that 2OGD-s44 enzyme unlikely catalyzes conversions of flavonoids (**Figure S11**). 2OGD-s44 belonged to a clade well supported by bootstrapping (98% bootstrap) with representatives from ferns, gymnosperms and angiosperms (**Figure 7B**). Each of the two *Azolla* genes assigned to the clade had a homologue from *A. caroliniana* (**Figure S12**). The clade’s angiosperm enzymes contained the *JASMONIC ACID OXIDASE (JOX)1-4* genes from Arabidopsis (**Figure 8B**), encoding JA-oxidases. The JOX were also the only Arabidopsis enzymes in the clade, suggesting that the clade represents enzymes accepting only a single substrate. Protein alignments revealed that the amino acids reported to interact with JA were conserved in the *Azolla* enzymes from this clade (Afi_v2_s44G002700.1 and Afi_v2_s11G001170.4) which further confirms them as very likely AfiJOX (**Figure S12**).

## Discussion

### The glycosylated trans-A triketide δ-lactone from insects is a semiochemical novelty

Insects are known for to recruit metabolic capabilities from bacteria and therefore are a rich source of allelopathic chemicals, compounds that mediate environmental signaling (Davis et al., 2013; Ferrari and Vavre, 2011). We have yet to reveal the ecological function of cornicinine and therefore cannot call it allelopathic. Its specific occurrence in the *N. cornicinina* species and its specific activity on *Azolla* ferns sharing the wetland habitat suggest that cornicinine is semiochemical. It is not volatile, however, and accumulates in the crane flies at levels much higher than would be expected from a pheromone. Its systematic association with adult *N. cornicina* collected through the insect wide range of distribution (**Figure S1C**) suggests that the crane fly synthesizes cornicinine with its own polyketide synthase (PKS). Recently, PKS were implicated in the biosynthesis of carminic acid, the red colorant from the cuticle of cochineal insects including *Dactylopius coccus* (Frandsen et al., 2018; Yang et al., 2021). PKS from animals including insects, however, have yet to be characterized (Frandsen et al., 2018).

Polyketide semiochemicals for which the biosynthesis pathway has been characterized thus far, have been synthesized by bacteria or fungi associated with insects. In some cases the microbes were insect defensive symbionts (Oliver and Perlman, 2020; van Moll et al., 2021). However, polyketides synthesized by microbes associated with insects were generally more complex than the comparatively small cornicinine aglycone with m/z 170. An example is the polyketide lagriamide of m/z 750 synthesized by the *Burkholderia* species associated with the beetle *Lagria villosa* (Flórez et al., 2018).

Cornicinine, a reduced triketide with a single glucose attached, resembles the simple polyketides synthesized by *Gerbera hybrida* plants, gerberin and parasorboside, identified as markers for the protection of the plants against oomycete fungi (Mascellani et al., 2022). Cornicinine has previously been extracted from the flowers of *Centaurea parviflora* that belong to the family of the Asteraceae as does *G. hybrida* (Belkacem et al., 2014). The PKS for the biosynthesis of *G. hybrida* triketides has been identified as well as the accessory enzymes for reduction of the pyrone which actually occurs before cyclization (Zhu et al., 2022). Accumulation of cornicinine in the crane fly could thus also result from its feeding behavior and that of its larvae. Identification of the PKS in the biosynthesis of cornicinine will reveal which organism synthesizes the compound in the future. The enzyme is of particular interest because a PKS synthesizing the R,R triketide lactone as in cornicinine has not been described: the PKS from antibiotic modules have proven very selective and difficult to engineer for a broader variety of substrates (Yin et al., 2003). PKS with novel properties have important applications to engineer novel polyketide drugs in pharmacology and (bio)pesticides in agriculture (Li et al., 2021). As such cornicinine could serve as the starting point for developing an *Azolla*-fern specific herbicide. Inactivity of the aglycone from cornicinine is consistent with previous results: polyketides require glycosylation for increased activity, uptake and transport, or stability (Mrudulakumari Vasudevan and Lee, 2020).

### Do trichomes lining *Azolla* leaf cavities mediate the response to cornicinine?

High expression of the LAR in the leaf-cavity preparations is linked to the trichomes since proanthocyanins accumulate there and it may be linked to the high JA-oxidase expression (**Figure 6, Figure 7**; Tran et al., 2020). In angiosperms, the link between JA elicitation and increased flavonoid accumulation is known and that between microbes inducing the JA-pathway and flavonoid accumulation is also well established (Albert et al., 2018; Chang et al., 2021). JA-control of glandular trichome differentiation and secondary metabolism is particularly well documented in tomato, but also found in artemisia (Ma et al., 2018; Xu et al., 2018). In contrast, bryophytes lack key enzymes of JA-Ile biosynthesis from the 12-oxo-phytodienoic acid precursor (OPDA). This includes *Marchantia polymorpha* that was reported to instead use OPDA-mediated signaling, thus not requiring JA-oxidase enzymes (Soriano et al., 2022). Consistently, the JA-oxidase clade supported with a bootstrap value of 98 (**Figure 7B**) did not contain sequences from the bryophytes and lycophytes; JA oxidation by JOX, therefore, evolved in the last common ancestor of ferns and angiosperms. A particularly interesting finding from the DOXC phylogeny (**Figure 7B, Figure S11**) was the position of the FLS/ANS clade of enzymes from the flavonoid biosynthesis as a sister clade to the JOX clade suggesting that the JOX and FLS/ANS evolved from an ancestor enzyme, by gene duplication, in the common ancestor of ferns and lycophytes, which may explain commonalities in their regulation.

### Cornicinine may function as an elicitor involving JA-metabolites

Many instances have been reported wherein semiochemicals from insects or plants alter bacteria physiology, yet in the present case the signal likely is host mediated because it only altered the cyanobiont, not free-living cyanobacteria. Also, leaf-cavities with characteristic trichomes develop in *Azolla* in the absence of the cyanobiont (**Figure 5A**).

The mechanism of host-control over the differentiation of Nostoc may involve the plant JA-pathway because of the high and specific expression of a JA-oxidase in host cells lining the akinetes when ferns were exposed to cornicinine and in megasporocarps. Accumulation of RNA encoding a glycolipid transferase and an allene oxidase in the leaf cavities of ferns on cornicinine suggested increased JA-synthesis and turnover into 12-OH-JA or 12-OH-JA-Ile (**File S1**). Given that methyl Jasmonate seemed ineffective in Azolla, the hydroxylated JA forms may be the active metabolite (De Vries et al., 2018). 12-OH-JA-Ile was recently shown to be an active JA form causing accumulation of anthocyanins in tomato and sorghum (Poudel et al., 2019)If the accumulation of active JA forms stretched to the whole leaf this would be consistent with cornicinine-induced leaf chlorosis (Jiang et al., 2014).

JA is a known player in plant immunity and its pathway may have been co-opted for symbiosis crosstalk in *Azolla*. Since the JA/SA pathway antagonism has been documented in bryophytes such as Marchantia, we expect both pathways to be active in the pteridophyte lineage and thus in *Azolla*. The core components of both pathways, the CRUL JA-receptor components COI1 and NPR1 were present in Marchantia; in addition, Marchantia reacted to necrotrophic and biotrophic pathogens with either JA/SA pathway in a manner similar to what is predicted from seed plants (Matsui et al., 2020). JA-receptors have yet to be characterized using the most advanced *Azolla* genome annotation (Afi_v2) released with this study, but they have been inferred by homology predictions in these ferns (de Vries et al., 2018).

Plants are known to perceive small molecules by way of CRUL complexes (Harper et al., 2021). Even simple metabolites such as quinone were shown to be sensed by CRUL (Laohavisit et al., 2020). The ominous accumulation of transcripts encoding several components of such system in the leaf cavities of cornicinine grown ferns but not in megasporocarps (**Figure 6D**) suggests that cornicinine may be sensed by a CRUL complex and thus may function as an elicitor. Elicitors from insects that trigger plant immunity have been characterized mostly from grazing and sucking insect pests but not crane flies (Jones et al., 2022; Santamaria et al., 2018). They are not generally volatile, they identified as peptides, fatty acid derivatives, for example fatty acid conjugated to glutamine or glutamate (FACS), or hydroxypropanoate esters of long-chain α, ω-diols. FACS accumulate at substantial levels because of their role in nitrogen assimilation in the insect gut (Yoshinaga et al., 2008). Responses to insect elicitors are specific for each system and stage (herbivory, oviposition), but were often associated with altered JA-accumulation. Cornicinine elicitation reduces nitrogen fixation because it induces akinete formation (**Figure 3**); reduced plant nitrogen may have evolved to reduce the fitness of the crane fly larvae which would presumably feed on the *Azolla* canopy once hatched.

### If JA mediates cornicinine elicitation via JOX, what is the fern response to cornicinine causing akinetes to form?

Transcripts encoding the sulfate transporter and the tetraspanin 8 accumulated in Azolla tissues where akinetes form. Sulfate or the lack of it has previously been shown to induce akinete formation, for example, in *Nostoc* ANTH a symbiotic strain known to colonize the roots of rice plants (Kyndiah and Rai, 2007; Wolk, 1965). Moreover, the sulfate transporter was listed in the repertoire of key genes specific for all symbiotic species of *Nostoc* (Warshan et al., 2018). Epiphytic colonization of ^33^S-labelled moss gametophytes showed furthermore that S-compounds are transferred to the *Nostoc punctiforme* from the feathermoss *Pleurozium schreberi* (Stuart et al., 2020). The structures of the shoot apex, the leaf cavity and the chamber of the indusium in *Azolla* are surrounded by hydrophobic envelopes, unlikely letting mineral nutrients pass from the surrounding water. The pore of the leaf cavity is adaxially oriented and water penetration is prevented by closure of the gap between the upper and lower leaf lobe. Nostoc is entirely dependent, therefore, on mineral supply from the fern throughout the life cycle of the symbiosis.

Electron microscopy revealed membrane vesicles (MV) surrounding Nostoc upon akinete formation in the megaspore indusium chamber from *A. microphylla* (Zheng et al., 2009). Images obtained after immunogold labeling demonstrate Nostoc cells fused with MV containing nucleic acids. Arabidopsis tetraspanin 8 knockout mutants were shown to secrete fewer extracellular vesicles than the wild types and such MV were found to contain small RNA that target fungal pathogens (Cai et al., 2018; Liu et al., 2020; Regente et al., 2017). The upregulation of tetraspannin 8 transcript in this study suggests that MV release is by the fern and facilitated by tetraspannin 8. Their content in nucleic acid is of particular interest because nucleic acids have been identified as a key in the maintenance of a phototrophic endosymbiosis: rRNA digestion products from the symbiont inhibit key transcriptional activity of the host which couples symbiont rRNA turnover with host vigor (Jenkins et al., 2021).

### No trace of ammonium transporter but sky-rocketing read numbers encoding the α-carbonic anhydrase and a SLAC channel in cells lining the leaf-cavity

Transcripts of AMT transporters or NOD26, known to transport ammonium/ammonia did not accumulate (**Figure S12**), in spite of predictions from other N_2_-fixating symbioses (Hwang et al., 2010). The pH surrounding symbiotic Nostoc was shown to be of importance in a symbiosis of peatmoss with *Nostoc muscorum* (Carrell et al., 2022). The pH of *Azolla* leaf cavities may be increased if by active Nostoc photosynthesis; it was reported to be 6.5 in leaves wherein Nostoc actively fixes N_2_. NH_3_ converted from the NH_4_ at this pH may not need a transport mechanism for uptake into the plant cell (Canini et al., 1992).

The very high accumulation of NRT1/PTR (Afi_v2_s47G001780.1) in addition to that of NRT3 (Afi_v2_s12G001250.1) is perplexing in the face of the complete absence of nitrate in the growth medium, and the low nitrate reductase transcripts in the sporophyte and absence in leaf cavities (**Figure 6A**). But nitrate may be synthesized via nitrogen oxide (NO) production from polyamines, hydroxylamine or arginine, the latter is synthesized in abundance. NO production and respiration was shown to be a pre-requisite for efficient N_2_-fixation in nodules from rhizobia (Valkov et al., 2020).

Teat cell studies suggest that they control gas exchange which would be of crucial importance to maintain CO_2_ and N_2_ in the leaf cavity (Veys et al., 2002, 2000, 1999). The sky rocketing levels of an α-carbonic anhydrase and a SLAC transcript in the LP profiles may thus stem from the leaf cavity pore, the opening of which may be dynamically controlled as in the case of stomata.

## Conclusion

Coordinated differentiation underlies the development of symbioses with filamentous cyanobacteria, including *Azolla*. A glycosylated triketide delta lactone, cornicinine, accumulates only in *N. cornicinina* crane flies that share the *Azolla* wetland habitat. Cornicinine targets the cyanobiont differentiation into akinete resting stages and thus inhibits N_2_-fixation and sexual reproduction of the *Azolla* symbioses. Cells lining the cyanobiont cavity exhibit transcriptional profiles consistent with cornicinine triggering plant elicitation. The results, including the Azfivs2 genome release, advance our understanding of the poorly studied but ecologically significant symbioses of seed-free plants with filamentous cyanobacteria.

## Materials and methods

### *Azolla* strains and growth conditions

The four *Azolla* species used in this study were *A. filiculoides* (Li et al., 2018), *A. pinnata* originating from the Botanical Gardens of Antwerp (Belgium), an *Azolla* species from the Anzali lagoon (Iran) which was phylogenetically analyzed but could not be assigned to any of the described *Azolla* species (Dijkhuizen et al., 2021) and an unknown *Azolla* species collected from the Botanical Gardens of Bordeaux (France). Adult *Azolla* sporophytes were grown in modified IRRI-medium as previously described (Brouwer et al., 2017) with a 16 h light period (100 µmol m^−2^ s^−1^) at 21°C.

### Preparation of *Azolla* spores for germination experiments

Spores for germination experiments were harvested in fall 2019 from mature mats of *A. filiculoides* by giving the plants a pressurized shower through a set of sieves (mesh sizes: 1000, 500 and 200 μm). Harvested spores were stored embedded in sludgy root debris at 4 °C. Shortly before use the sludge was diluted with water and agitated in a wide container. Distinguishably yellow-colored clumps of megasporocarps, held together by the glochidia of the massulae, could then be hand-picked from the shallow water and used.

### *Nephrotoma cornicina*, isolation, bioassay and structural analyses of cornicinine

The thousands *Nephrotoma cornicina* (Linnaeus, 1758) (Tipulidae, Diptera) used for the initial identification of cornicinine were collected in the surroundings of Louvain-la-Neuve (Belgium). Entomologists from various parts of the world kindly provided material from their country (see figure 1 - figure supplement 1C). Bioassay, isolation procedure and structural elucidation of cornicinine has been described in patent EP1697392A1 (Mathieu et al., 2005). Briefly, bioassays of fern fronds in liquid medium were carried out with 4 μg ml^-1^ of dry *N. cornicina* powder. For structural analyses, aqueous extract from *N. cornicina* (about 10,000 adults) was fractioned on a Sephadex G-10 column and assayed for bioactivity. 35 mg of a pure compound could be isolated from the bioactive fractions with preparative HPLC on a C18 column. APCI/HREI mass spectrometry revealed the compound had a sugar moiety and the mass of the isolated aglycone corresponded to C_9_H_14_O_3_. NMR experiments (INADEQUATE, HSQC, HMBC, COSY, ROESY and NOE) followed by structural analyses revealed a novel glycosylated triketide δ-lactone which was called cornicinine (C_15_H_24_O_8_). Cornicinine could have three possible isomeric configurations (cis, trans-A and trans-B) but the trans-configuration fitted best with the NMR-data.

### Chemical synthesis and characterization of cornicinine stereoisomers and their aglycone precursors is described in Method S1

#### Cornicinine assays on *Azolla*

The synthesized compounds were tested by putting ±3 mm *Azolla* shoot tips in 1.5 ml IRRI-medium in a 24-well plate and supplementing 500 nM of each compound dissolved in water. 1 mM KNO_3_ was added to the IRRI-medium to test the effect of nitrate and cornicinine together on *Azolla*. To test the effect of cornicinine on germinating sporelings, clumps of ±50 megasporocarps were inoculated in 1.5 ml water supplemented with 500 nM cornicinine. Buoyant sporelings surfaced after 10 days and were transferred to fresh IRRI-medium with 500 nM cornicinine during further development.

#### Cornicinine assays on *Arabidopsis thaliana* and free-living filamentous cyanobacteria

*A. thaliana* Col-0 seeds were sterilized for 3 h by chlorine gas vapor and sown on ½ MS medium including vitamins pH 5.8 with 0.8% agarose and 500 nM cornicinine. The seeds were stratified for 2 days at 4 °C and moved to a 16 h light period (100 µmol m^−2^ s^−1^) at 21°C. Anabaena sp. PCC 7210 was inoculated in BG-11 medium with 500 nM cornicinine and grown under constant light (25 µmol m^−2^ s^−1^­) at 30 °C. Growth was tracked by measuring OD_665_ of methanolic extracts and the formula: chlorophyll (mg/ml) = 13.45 * OD_655_ * dilution factor. Akinete induction was tested on free-living cyanobacteria donated by Henk Bolhuis (NIOZ-Texel, The Netherlands), incubating them in BG-11 medium in the presence of 500 nM cornicinine.

#### Microscopy

*A. N. azollae* was visualized by squeezing the outermost tip of an *Azolla* branch between two glass slides with a drop of demineralized water. A Zeiss Axio Zoom.V16 microscope with a Zeiss Axiocam 506 color camera, Zeiss CL 9000 LED lights and a Zeiss HXP 200C fluorescence lamp with standard Zeiss RFP filter set 63HE (excitation 572 nm, emission 629 nm) was used for imaging. Images were Z-stacked with Helicon Focus 7 software in default settings (depth map, radius 8, smoothing 4). The same set-up was also used to image sporelings, leaf cavities and free-living filamentous cyanobacteria.

#### Leaf-cavity isolations from *A. filiculoides* and sequencing of their polyA-enriched RNA

Leaf cavities were isolated from *A. filiculoides* as described before with slight modifications (Peters et al., 1978; Uheda, 1986). Briefly, about 3 g of *Azolla* was prepared by removing roots and rinsing with 0.1% v/v Triton X-100 and demineralized water. Cleaned sporophytes were submerged in enzyme solution (0.5 M mannitol with 2% w/v cellulase, 1% w/v macerozyme, 0.1% w/v pectolyase, 1% w/v PVP and 10 mM DTT) and vacuum infiltrated for 10 min at 0.6 bar before incubation for 19 h at 30 °C with gentle agitation. Leaf cavities were released by washing the digested sporophytes with 0.5 M mannitol through a mesh. The flow-through was left to settle for 10-30 min after which the sunk leaf cavities were manually collected and washed in PBS before snap freezing.

The experiment was set up so as generate three biological replicates to compare RNA extracted from sporophytes with that of isolated leaf cavities, and to compare RNA in leaf cavities isolated from ferns grown with and without 500 nM cornicinine for 6 days. Care was taken to snap-freeze the ferns and isolated leaf cavities 2-3 hours into the light cycle of the diel rhythm. Total RNA from ±80 isolated leaf cavities was extracted with the RNeasy Micro Kit (Qiagen, Germany). Total RNA from 50 mg FW sporophytes was isolated with the Spectrum Plant Total RNA Kit (Sigma-Aldrich) applying protocol B. Total RNA was treated with DNase I (Thermo Fisher Scientific, Waltham, Massachusetts, USA) for 1 h at 37 °C after which the reaction was stopped by adding two mM EDTA and incubation for 10 min at 65°C. The reactions were cleaned with the RNeasy MinElute Cleanup Kit (Qiagen). Poly-A tail enriched cDNA libraries were prepared using the SMART-Seq HT Kit (Takara Bio, Japan), quality checked using the TapeStation DNA ScreenTape (Agilent Technologies, Santa Clara, California, USA) then sequenced on a half lane NovaSeq 6000 using the paired-end 2x50 cycle chemistry (Ilumina, San Diego, California, USA). Data is deposited under accession nr. (provided upon acceptance of the manuscript).

#### Dual RNA-Sequencing of far-red light grown sporophytes, micro and megasporocarps

*A. filiculoides* ferns were grown on light with a far-red component to induce sporulation as described in Dijkhuizen et al., 2021. Micro and megasporocarps were manually picked from the sporulating ferns during a period of two h, two h into the light period, snap frozen along with the sporophytes collected two h into the light period. Material was sampled from independent cultures so as to obtain three independent biological replicates for each of the megasporocarp, microsporocarp and sporophyte samples. RNA was extracted, then libraries synthesized and dual RNA sequenced as described in Dijkhuizen et al., 2021. Data from this experiment is deposited under accession nr. (provided upon acceptance of the manuscript).

### Sequencing, assembly and annotation of the *A. filiculoides* genome version 2 (Afi_v2) is described in Method S2

#### Analysis of the differential mRNA accumulation in leaf-cavity preparations

After demultiplexing, quality filtering and trimming of the sequencing primers away from the reads, approximatively paired reads per sample were aligned using the STAR aligner with default settings to the concatenated genome assemblies of the *A. filiculoides* nucleus Afi_v2, its chloroplast and *N. azollae*, extracting read counts for Afi_v2 only.

Read counts for the Afi_v2 gene models (predominant splice versions only) were normalized as reads per million, except for leaf-cavity profiling. For the later, normalization was to the sum of counts from the 1100 most-expressed genes in each sample because of the large difference in the sensitivity of the assay when comparing sporophytes with leaf cavity. For statistical analyses of differential gene expression with DESeq2 (Love et al., 2014), the genes with no expression in all leaf cavity samples were removed from the analyses. In addition, the sample leaf cavity 2 from sporophytes grown on cornicinine was removed from the analysis because of its large contamination with sporophyte RNA.

#### Phylogenetic analysis of genes encoding 2-oxoglutarate-dependent dioxygenases (2OGD)

Protein sequences of 2-oxoglutarate-dependent dioxygenases in the two *A. filiculoides* genome assemblies and annotations were identified by local BLAST using as query: automatically annotated as Azfi 2OGD genes by Mercator (Lohse et al., 2014). These were compared to functionally characterized DOXC-genes from seed plants (Kawai et al., 2014) and bryophytes (Li et al., 2020). Phylogenies were created of these sequences in the context of a DOXC orthogroup obtained from the 1kp orthogroup database (Ka-Shu Wong et al., 2019). The orthogroup was sub-sampled and sequences were aligned with MAFFT-einsi (Katoh et al., 2019), and then trimmed using trimAL (Capella-Gutierrez et al., 2009). The phylogeny was computed with IQ-tree (Nguyen et al., 2015) with 200 bootstraps. Bootstrap support was calculated as transfer bootstraps (Lemoine et al., 2018). A subset of the phylogeny containing JOX, ANS and FLS clades was re-computed similarly. Both trees were annotated in iTOL (Letunic and Bork, 2019) and Inkscape. Code and data for the phylogeny are available at https://github.com/lauralwd/2OGD_phylogeny.

## Supporting information

Read count tables for fern genes expressed in the leaf cavities and in megasporocarps

## Acknowledgements

We would like to thank Pjotr Oosterbroek for taxonomic assignments and for his help in contacting entomologists who provided *Nephrotoma cornicina* from various parts of the world. We thank Pasquale Ciliberti from Naturalis Biodiversity Center (Leiden, Netherlands) and Dr. Henk Bolhuis from Royal Netherlands Institute for Sea Research (Texel, Netherlands) for sharing *N. cornicina* crane fly specimens, and free-living filamentous cyanobacteria respectively. We further would like to thank Nils Stein for hosting the HiC work and Ines Walde for her technical help on the Hi-C library preparations and sequencing at the IPK (Seeland, Germany).

**Accessions** of sequencing data and genome assembly and annotation will be provided upon acceptance of the manuscript

## Supporting Information

### Supporting Methods

#### Method S1. Chemical synthesis and characterization of cornicinine stereoisomers and their aglycone precursors

Freshly distillated dibutylboron trifluoromethanesulfonate (Bu_2_BOTf, 8.6 mL, 34 mmol, 2 eq) in diethylether (Et_2_O, 16 mL) was slowly added to a solution of propionyl oxazolidinone precursor (4 g, 17 mmol) in Et_2_O (52 mL) at 0 °C (**Figure S2A**, step a). N,N-Diisopropylethyleneamine (DIPEA, 3.4 mL, 20 mmol, 1.15 eq) was then added at such a rate as to keep the internal temperature below 2°C. Once the addition was complete, the mixture was cooled to −78 °C before freshly distilled propionaldehyde (1.5 mL, 21 mmol, 1.25 eq) in Et_2_O (20 mL) was introduced. The resulting mixture was stirred for 30 min at −78°C, and then for 1h at 0°C. The reaction was quenched at −78°C with tartaric acid (9 g). The resulting mixture was stirred for 2 h at room temperature. Water (50 ml) was then added and the aqueous layer was extracted with ether (2x 25 mL). The combined organic layers were washed with saturated NaHCO_3_ (2x 25mL). The organic layer was then transferred to a round bottom flask, cooled to 0°C, so as to obtain a 3:1 mixture of MeOH/ 30% H_2_O_2_. After 30 min at room temperature, the solution was extracted with ether (2x 40mL), and washed with saturated NaHCO_3_ (40 mL), and brine (40 mL). The volatiles were removed under vacuum and the product was used directly without further purification.

Freshly distillated propionic anhydride (4.4 mL, 34 mmol, 2eq), followed by triethylamine (4.7 mL, 34 mmol, 2eq) and 4-dimethylaminopyridine (DMAP, 0.1 eq) were added to a solution of the crude alcohol (17 mmol, 1 eq) in dichloromethane (DCM, 16 mL) (**Figure S2A**, step b). The resulting mixture was stirred 2h, then washed with 1M HCl (2x 10 mL), H_2_O (2x 10 mL), saturated aqueous solution of NaHCO_3_ (2x 10 mL), and brine (2x 10 mL). The organic phase was dried over Na_2_SO_4_, filtered and then the volatiles were removed under vacuum. The residue was purified by silica gel chromatography (PE:EtOAC, 95:5 to 60:40) to afford the ketone product with 52% yield over two steps (**Figure S2A**, step a-b). NMR analyses yielded data in agreement with literature (Hinterding et al., 2001).

To a solution of ketone (0.520 mg, 1.5 mmol, 1 eq) in tetrahydrofuran (THF, 12 mL) was added dropwise a solution of potassium bis(trimethylsilyl)amide (KHMDS) in THF (4.5 mL, 4.5 mmol, 3 eq, 1M) at −78°C (**Figure S2A**, step c). The resulting mixture was stirred 1 h at −78°C, then quenched with a mixture of NH_4_Cl/MeOH/H_2_O (1:1:1 v/v/v, 30 mL) and warmed to room temperature. Ethylacetate (30 ml) and water (10 ml) were added and the layers separated. The organic phase contained the chiral auxiliary that was isolated for recycling. The basic aqueous phase was acidified to pH 2-3 with 0.25 M HCl, and then extracted with DCM (3x 50 mL). The combined organic layers were dried over Na_2_SO_4_, filtered and then the volatiles were removed under vacuum. The residue was purified by silica-gel chromatography (PE:EtOAC, 95:5 to 80:20) to afford the lactone product with 71% yield (**Figure S2B**, lactone trans-A). The product was further characterized by NMR. 1H NMR (300 MHz, CDCl3): δ 4.35 (ddd, J = 10.7, 7.9, 2.9 Hz, 1H), 3.55 (q, J = 6.6 Hz, 1H), 2.35 (dq, J = 10.6, 7.2 Hz, 1H), 1.96 (dqd, J = 14.8, 7.4, 3.0 Hz, 1H), 1.72 (dq, J = 14.6, 7.5 Hz, 3H), 1.38 (d, J = 6.7 Hz, 3H), 1.22 (t, J = 7.2 Hz, 4H), 1.16 – 1.06 (m, 4H). 13C NMR (75 MHz, CDCl3): δ 204.92, 170.14, 80.26, 50.19, 45.91, 25.04, 12.02, 8.71, 8.01. ES HRMS (m/z): Calculated for C9H13O3 (M-H): 169.08592; found: 169.08537

To a solution of lactone trans-A (0.110 mg, 0.65 mmol, 1 eq) in dimethylformamide (DMF, 2 mL), tetra-O-acetyl-β-glucosyl bromide (938 mg, 2.28 mmol, 3.5 eq) and Cs_2_CO_3_ (743 mg, 2.28 mmol, 3.5 eq) were added, at room temperature and protected from light (**Figure S2B**, step d). After 3h, H_2_O (10 mL) and DCM (10 mL) were added and the layers separated. The aqueous phase was extracted with DCM (2x 10 mL). The combined organic layers were washed with brine (3x 10 mL), and then dried over MgSO_4_, filtered and then volatiles removed under vacuum. The residue was purified by silica gel chromatography (PE:EtOAC, 50:50 to 0:100) to afford the product with 68% yield (**Figure S2B**, after step d), which was analyzed by NMR. 1H NMR (300 MHz, CDCl3): δ 5.27 (t, J = 9.1 Hz, 1H), 5.23 – 5.11 (m, 2H), 5.10 – 5.04 (m, 1H), 4.25 – 4.02 (m, 4H), 3.81 (ddd, J = 10.0, 5.2, 2.5 Hz, 1H), 2.87 (d, J = 6.7 Hz, 1H), 2.67 – 2.51 (m, 1H), 2.11 – 1.97 (m, 12H), 1.89 – 1.80 (m, 1H), 1.78 (s, 3H), 1.68 – 1.53 (m, 1H), 1.27 (d, J = 7.0 Hz, 3H), 0.98 (t, J = 7.4 Hz, 3H). 13C NMR (75 MHz, CDCl3): δ 170.45, 170.21, 169.39, 169.06, 166.28, 164.52, 136.05, 127.32, 108.49, 96.20, 82.46, 72.44, 72.31, 71.08, 68.04, 61.97, 32.46, 26.52, 20.76, 20.69, 20.62, 17.63, 10.20, 9.43. ES HRMS (m/z): Calculated for C23H32O1223Na (M+Na): 523.17860; found: 523.17878.

The same experiment, as above, was performed starting from lactone trans-B (0.102 mg) to afford the product with 65% yield (**Figure S2C**, step d) 1H NMR (300 MHz, CDCl3): δ 5.32 – 5.06 (m, 4H), 4.87 (d, J = 7.3 Hz, 1H), 4.23 – 4.16 (m, 2H), 4.15 – 4.02 (m, 1H), 3.76 (ddd, J = 10.0, 4.7, 3.3 Hz, 1H), 2.63 – 2.49 (m, 1H), 2.13 – 1.99 (m, 16H), 1.80 (d, J = 1.3 Hz, 4H), 1.74 – 1.57 (m, 3H), 1.27 (d, J = 6.9 Hz, 3H), 1.11 (dd, J = 11.1, 7.2 Hz, 2H), 1.03 (t, J = 7.4 Hz, 3H). 13C NMR (75 MHz, CDCl3): δ 170.37, 170.19, 169.35, 169.07, 166.50, 166.20, 110.33, 98.73, 82.48, 72.41, 72.36, 71.08, 68.00, 61.80, 35.37, 26.25, 20.68, 20.62, 20.58, 16.58, 9.94, 9.40. ES HRMS (m/z): Calculated for C23H32O1223Na (M+Na): 523.17860; found: 523.17893.

To a solution of protected sugar trans-A (140 mg, 0.28 mmol, 1 eq) in MeOH (3 mL), sodium methoxide (MeONa, 6 mg) was added (**Figure S2B,** step e). After 3 h, IR20-amberlite (H+) was added to the resulting mixture. The resulting mixture was stirred for 3 h and then filtered. The volatiles were removed under vacuum. The residue was purified by silica gel chromatography (PE:EtOAC, 70:30 to 0:100) to afford the pure product with 81% yield (**Figure S2B**, after step e). The product was characterized by NMR. 1H NMR (300 MHz, DMSO): δ 5.39 (s, 1H), 5.10 (d, J = 20.4 Hz, 2H), 4.82 (d, J = 7.3 Hz, 1H), 4.56 (s, 1H), 4.10 (ddd, J = 8.0, 5.7, 1.7 Hz, 1H), 3.67 (d, J = 11.7 Hz, 1H), 3.53 – 3.40 (m, 1H), 3.30 – 3.07 (m, 5H), 2.79 (q, J = 6.8 Hz, 1H), 1.68 (s, 3H), 1.75 – 1.48 (m, 5H), 1.25 (d, J = 6.9 Hz, 3H), 0.89 (t, J = 7.4 Hz, 3H). 13C NMR (75 MHz, DMSO): δ 168.47, 166.29, 104.17, 99.82, 82.41, 77.80, 76.86, 73.80, 70.04, 61.12, 49.06, 33.06, 26.32, 18.52, 10.58, 9.69. ES HRMS (m/z): Calculated for C15H24O823Na (M+Na): 355.13634; found: 355.13628.

The same experiment, as above, was performed starting from protected sugar trans-B (0.120 mg) to afford the product with 75% yield (**Figure S2C**, step e). 1H NMR (300 MHz, DMSO): δ 5.61 – 4.98 (m, 3H), 4.83 (d, J = 7.3 Hz, 1H), 4.51 (s, 1H), 4.11 (ddd, J = 7.8, 6.0, 1.4 Hz, 1H), 3.62 (dd, J = 11.8, 2.0 Hz, 1H), 3.45 (dd, J = 11.6, 5.4 Hz, 1H), 3.28 – 3.09 (m, 4H), 2.80 (q, J = 6.8 Hz, 1H), 1.65 (s, 3H), 1.73 – 1.53 (m, 2H), 1.22 (d, J = 7.0 Hz, 3H), 0.88 (t, J = 7.4 Hz, 3H)13C NMR (75 MHz, DMSO): δ 167.86, 166.17, 103.49, 98.69, 82.20, 77.57, 76.84, 73.58, 69.92, 60.96, 31.59, 26.41, 18.77, 10.63, 9.59.

#### Method S2. Sequencing, assembly and annotation of the *A. filiculoides* genome version 2 (Afi_v2)

To improve the first genome assembly (4666 scaffolds), *A. filiculoides* PacBio RSII sequencing data from Li et al., 2018 was processed anew. PacBio RSII reads were corrected, trimmed and assembled with Canu (Koren et al., 2017) in 4632 scaffolds. The assembly was then polished with Quiver (Chin et al., 2013) and Pilon (Walker et al., 2014). In order to reduce fragmentation, we implemented optical mapping. We grew *A. filiculoides* without cyanobacteria under sterile conditions, extracted nuclei as described in Dijkhuizen et al., 2018, extracted high molecular weight DNA (above 150kb) and ran Bionano Genomics chips once this DNA was labelled as per the manufacturer’s instructions. With the optical maps, the new *A. filiculoides* genome assembly was reorganized into 4422 scaffolds.

To further reduce fragmentation, we resorted to incorporating tethered chromosome conformation capture sequencing (TCC) . The TCC library was prepared essentially as described previously (Himmelbach et al., 2018) from 2.4 g of the same fresh plant material as for the optical mapping. The library was sequenced using a HiSeq2500 (Illumina Inc., San Diego, CA, USA). TCC data was then used to correct and improve the optically mapped assembly (Mascher et al., 2017). The final Afi_v2 assembly contains 3585 scaffolds totaling 579 Mbp with an N50 of 4 Mbp and L50 of 35. The assembly is deposited under accession nr. (upon acceptance of the manuscript). Afi_v2 is shorter than the 622.6 Mb, but its N50 is much improved over the N50 of 965kb, compared to the first assembly (Li et al., 2018).

To improve on the first *A. filiculoides* annotation, we included existing unstranded RNA sequencing data (Brouwer et al., 2017; Vries et al., 2016) and more recent stranded and dual RNA sequencing data (Dijkhuizen et al., 2021). RNA-sequencing data were mapped to the Afi_v2 assembly with STAR (Dobin et al., 2013) and gene predictions were made with GeMoMa (Keilwagen et al., 2019). The Afi_v2 annotation is deposited under sequence accession number (upon acceptance of the manuscript). We used a gene predominant splice form for read counting. For comparison, our searches for 2-oxoglutarate-dependent dioxygenase enzymes resulted in 22 gene models predicted for Azfi_vs1 but 29 for Afi_v2.

## Supporting Figures

**Figure S1.**
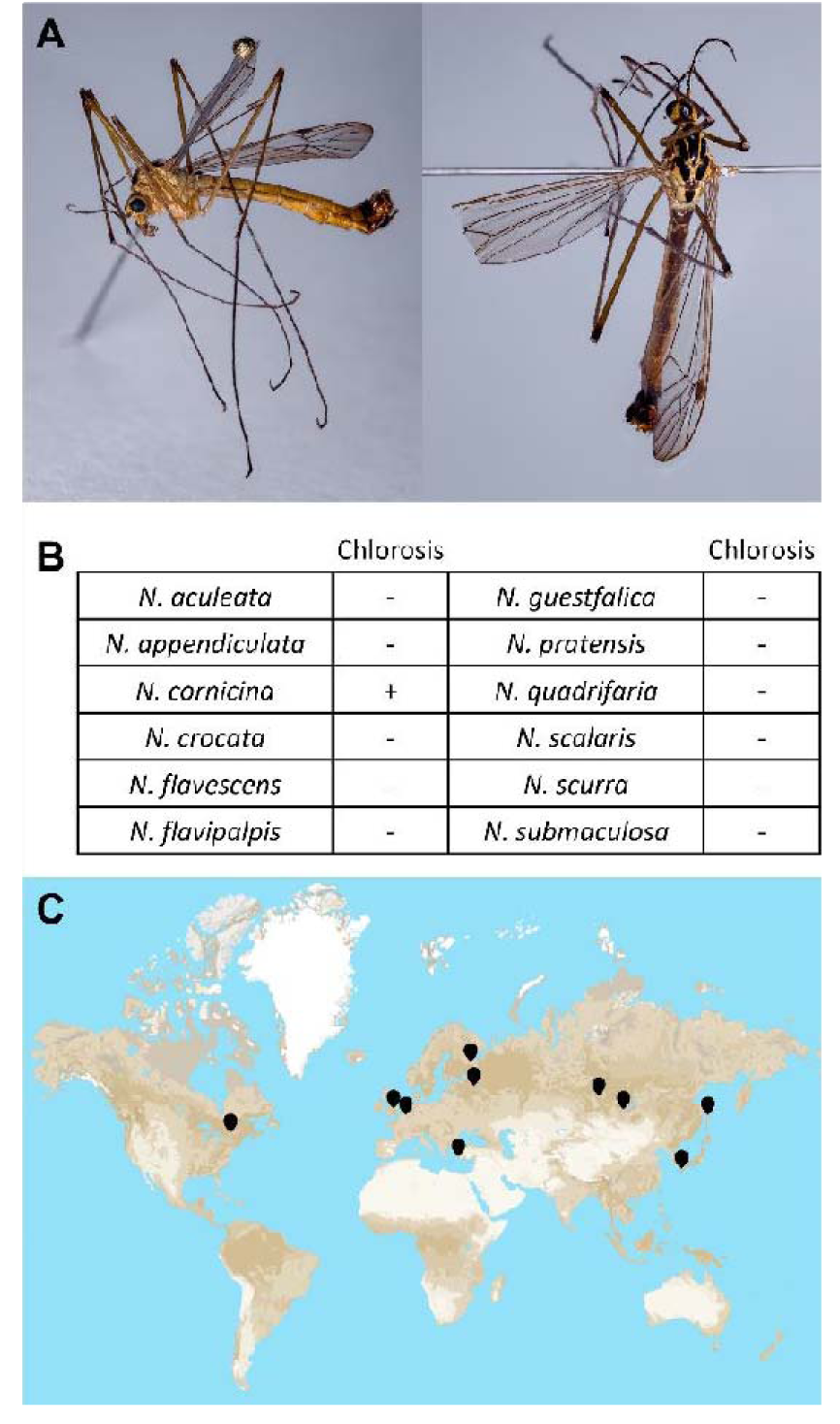
Specificity of the chlorosis induced by *N. cornicina*. (**A**) *N. cornicina* specimen from the collection of Naturalis Biodiversity Center (Leiden, Netherlands) collected in 1999 in the city of Nijkerk (Netherlands). (**B**) *Nephrotoma* species tested for their ability to induce chlorosis on mats of *A. filiculoides*. -, no chlorosis; +, chlorosis. (**C**) Collection sites of the specimens of *N. cornicina* tested in this study, all of which induced chlorosis.

**Figure S2.**
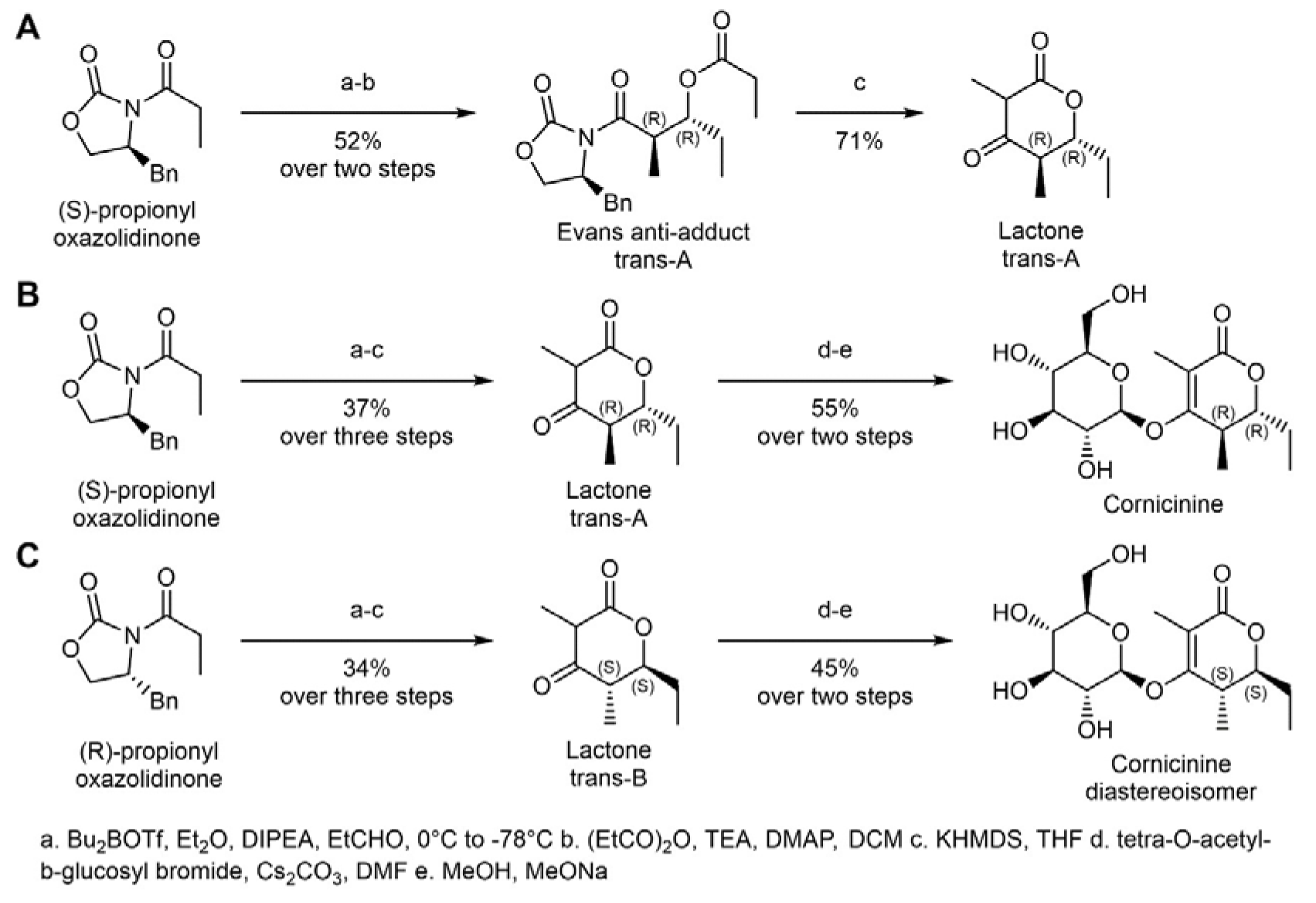
Chemical synthesis of cornicinine, its diastereoisomer, and their aglycones. (**A**) Starting from the commercial propionyl oxazolidinone, the synthesis of the lactone was achieved in three steps (a-c) using the conditions to provide the Evans antiadduct after step b. (**B**) and (**C**), The resulting alcohol from step b was immediately acetylated using propionic anhydride. The intramolecular lactonization was achieved using an excess of KHMDS at −78°C to provide the trans A (R,R) & trans B (S,S)-lactones with 37% and 34% yield, respectively after step c. The o-glycosylation with the tetra-O-acetyl-b-glucosyl bromide under basic conditions (Cs_2_CO_3_) provided the (R,R)-lactone o-glucosyl with 68% yield, and 65% yield for the (S;S)-lactone O-glucosyl in step d. Under basic methanolic conditions, the cornicinine product was obtained with 81% yield in step e. Under the same conditions, the diastereoisomer was obtained with 69% yield in step e. The overall yields over the steps a-c and d-e were as indicated.

**Figure S3.**
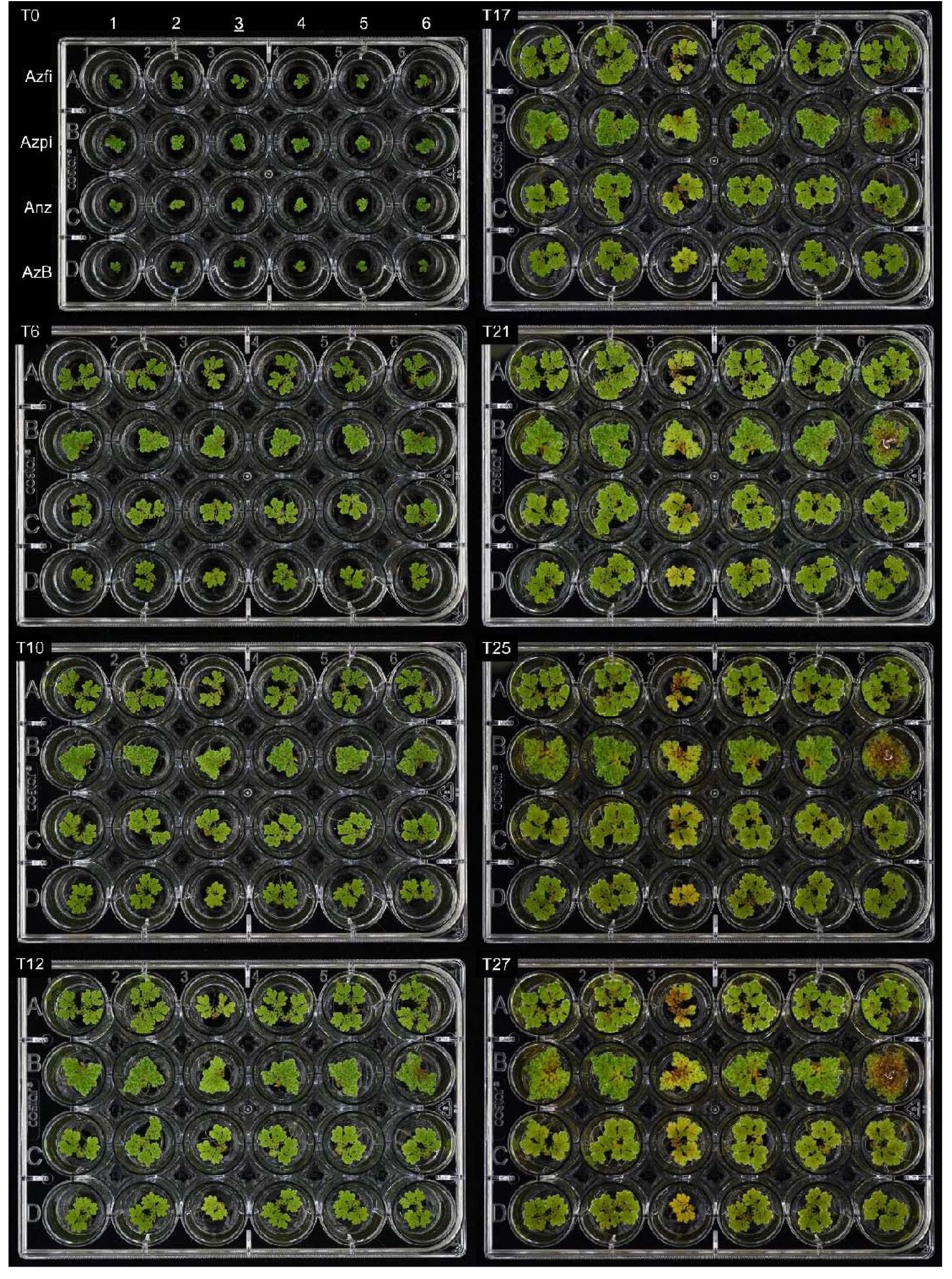
*Azolla* species grown for 27 days on 500 nM of the compounds from Figure 2. Azfi: *A. filiculoides*; Azpi: *A. pinnata*; Anz: *Azolla* species from Anzali (Iran); AzB: *Azolla* species from Bordeaux (France).

**Figure S4.**
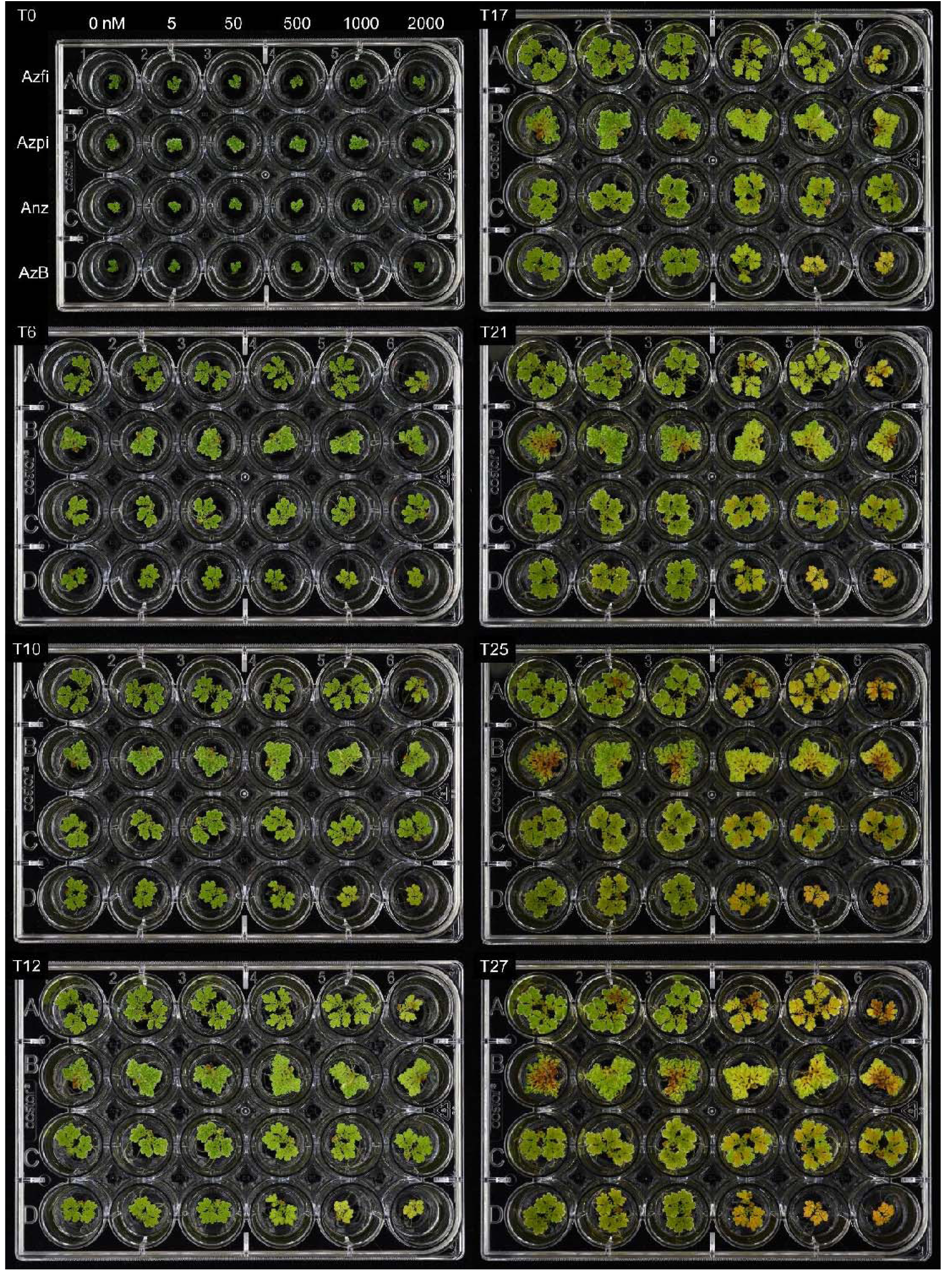
*Azolla* species grown for 27 days on a concentration gradient ranging from 0-2000 nM of the bioactive trans-A diastereoisomer cornicinine, <u>3</u> from Figure 2. Azfi: *A. filiculoides*; Azpi: *A. pinnata*; Anz: *Azolla* species from Anzali (Iran); AzB: *Azolla* species from Bordeaux (France).

**Figure S5.**
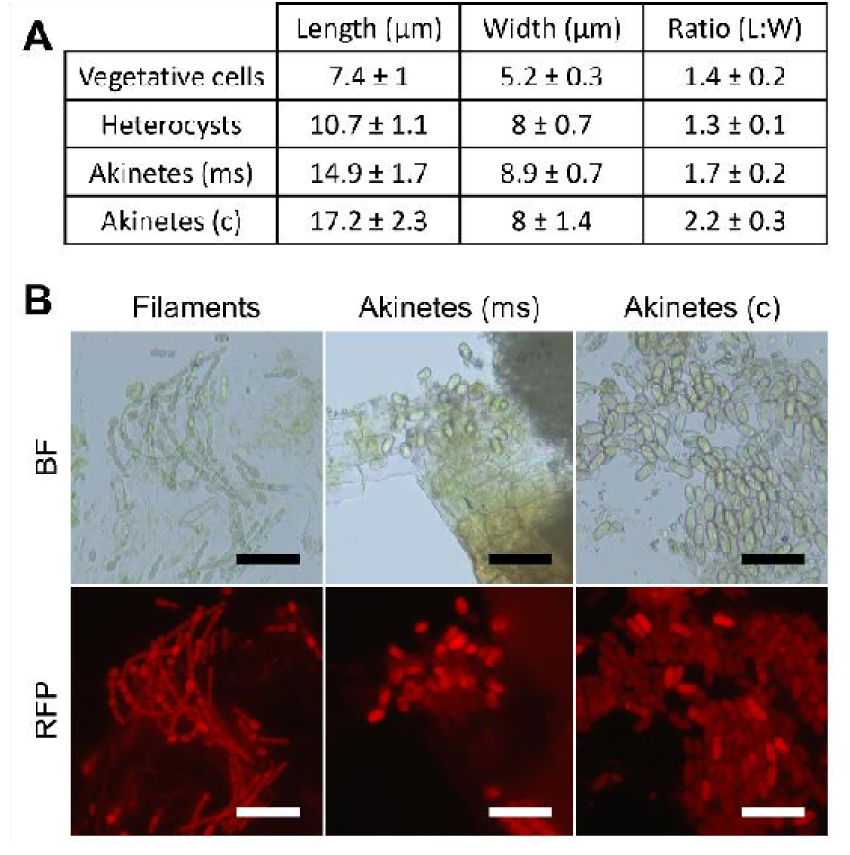
Size and morphology of the different stages of *N. azollae*. (**A**) Length and width of vegetative cells, heterocysts, akinetes in the megasporocarp (ms) and akinete-like cells induced by cornicinine (c). (**B**) From left to right, typical vegetative cells with heterocysts, akinetes in the megasporocarp (ms) and akinete-like cells induced by cornicinine (c) of *N. azollae*. BF: bright-field; RFP: red fluorescent protein settings. Scale bars correspond to 50 μm.

**Figure S6.**
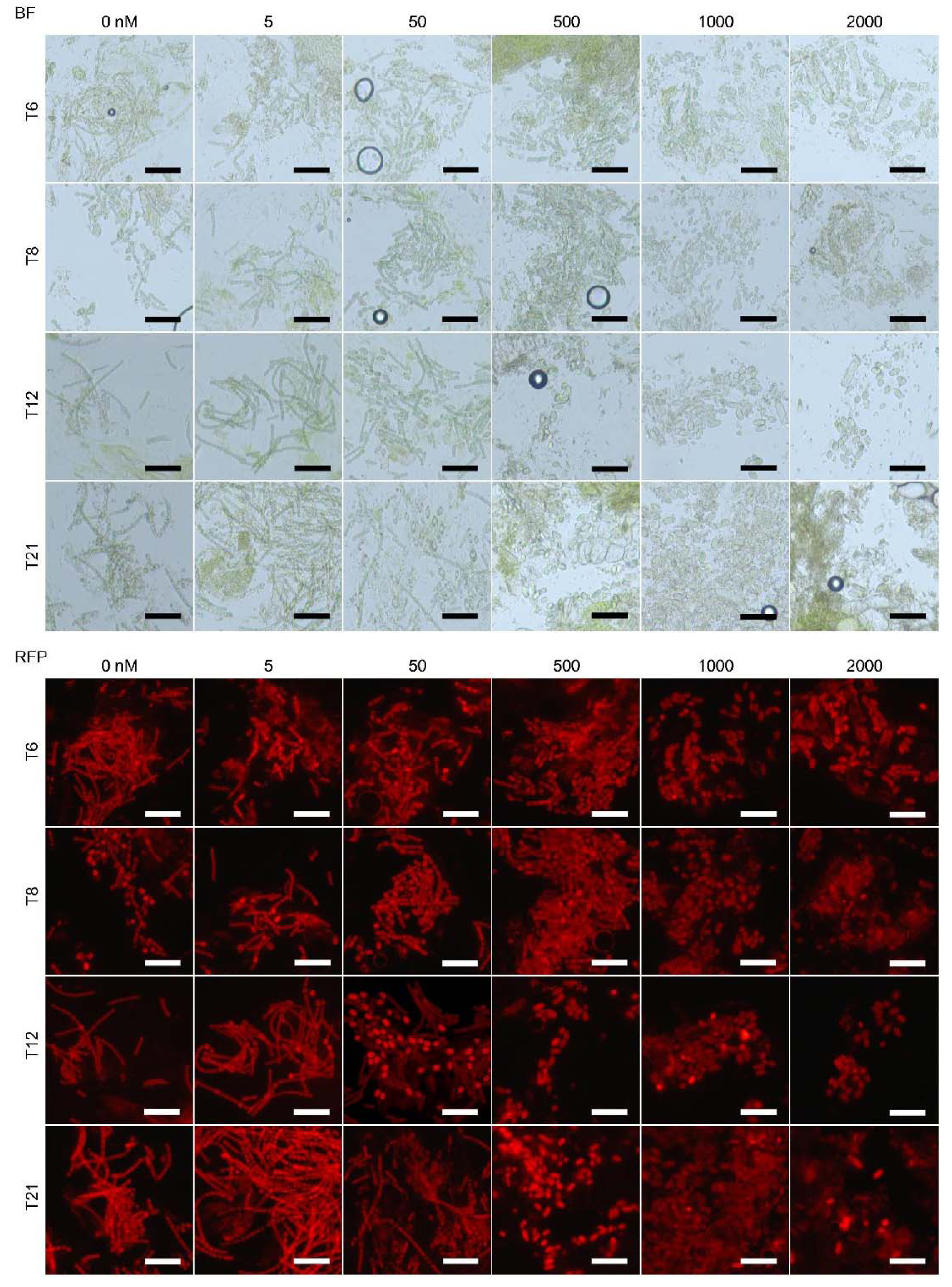
*N. azollae* development inside *A. filiculoides* treated with a concentration gradient from 0-2000 nM of cornicinine, <u>3</u> from Figure 2. BF: bright-field; RFP: red fluorescent protein settings. Scale bars correspond to 50 μm.

**Figure S7.**
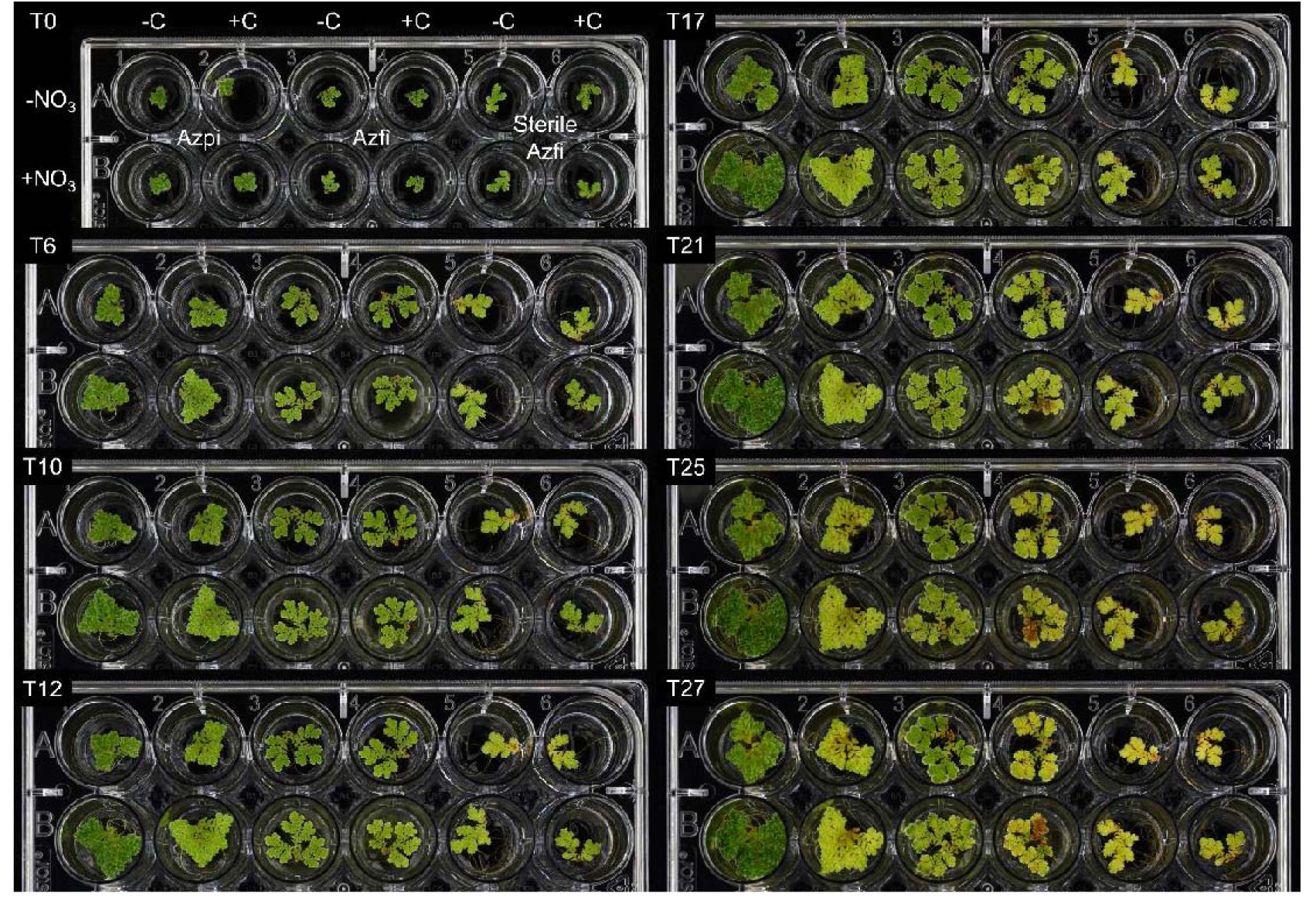
*Azolla* species grown for 27 days on medium supplemented with(out) 500 nM cornicinine (-/+C) and 1 mM KNO_3_ (-/+NO_3_). Azfi: *A. filiculoides*; Azpi: *A*.

**Figure S8.**
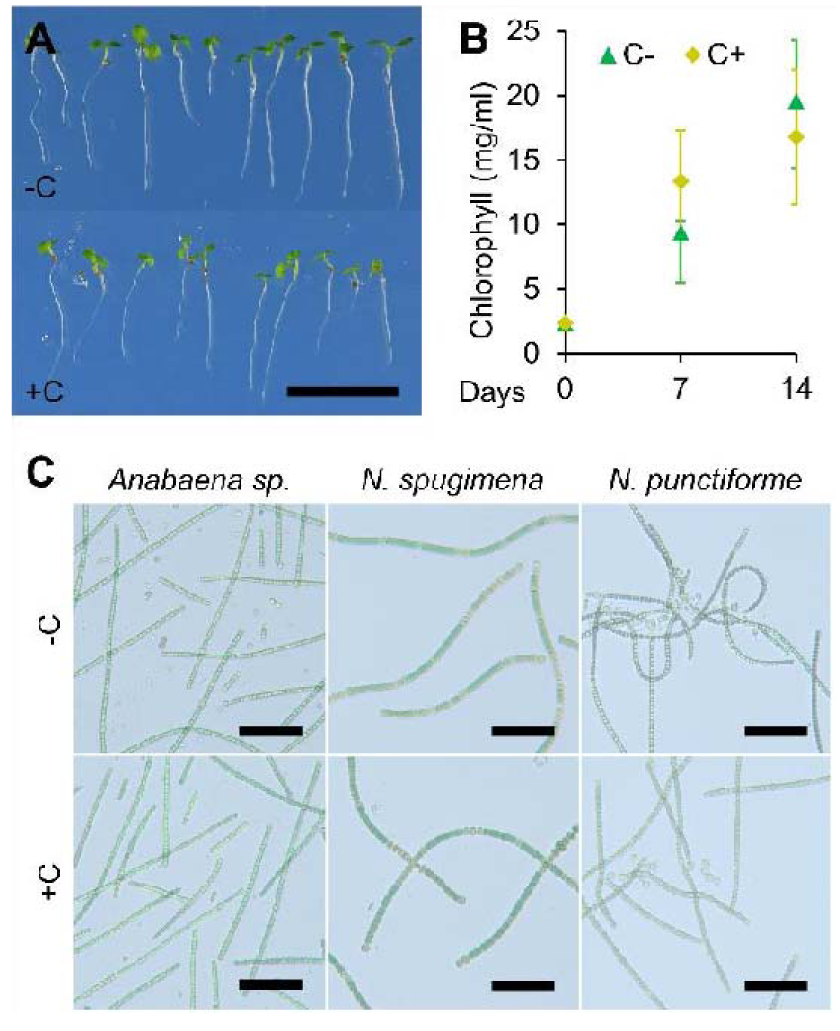
Effect of cornicinine on *Arabidopsis thaliana* and free-living filamentous cyanobacteria. (**A**) Effect of 500 nM cornicinine (-/+C) on germination and development of model plant *Arabidopsis thaliana* after seven days. (**B**) Effect of 500 nM cornicinine in the medium on the growth of the filamentous cyanobacterium Anabaena sp. PCC 7210 during 14 days. (**C**) Effect of 500 nM cornicinine (-/+C) in the medium of free-living filamentous cyanobacteria from the collection of Royal Netherlands Institute for Sea Research (Texel, Netherlands) after three weeks. All species are isolates from sediments on the island of Schiermonnikoog (Netherlands): Anabaena sp. (CCY1406), *Nodularia spumigena* (CCY1407) and *Nostoc punctiforme* (CCY1588).

**Figure S9.**
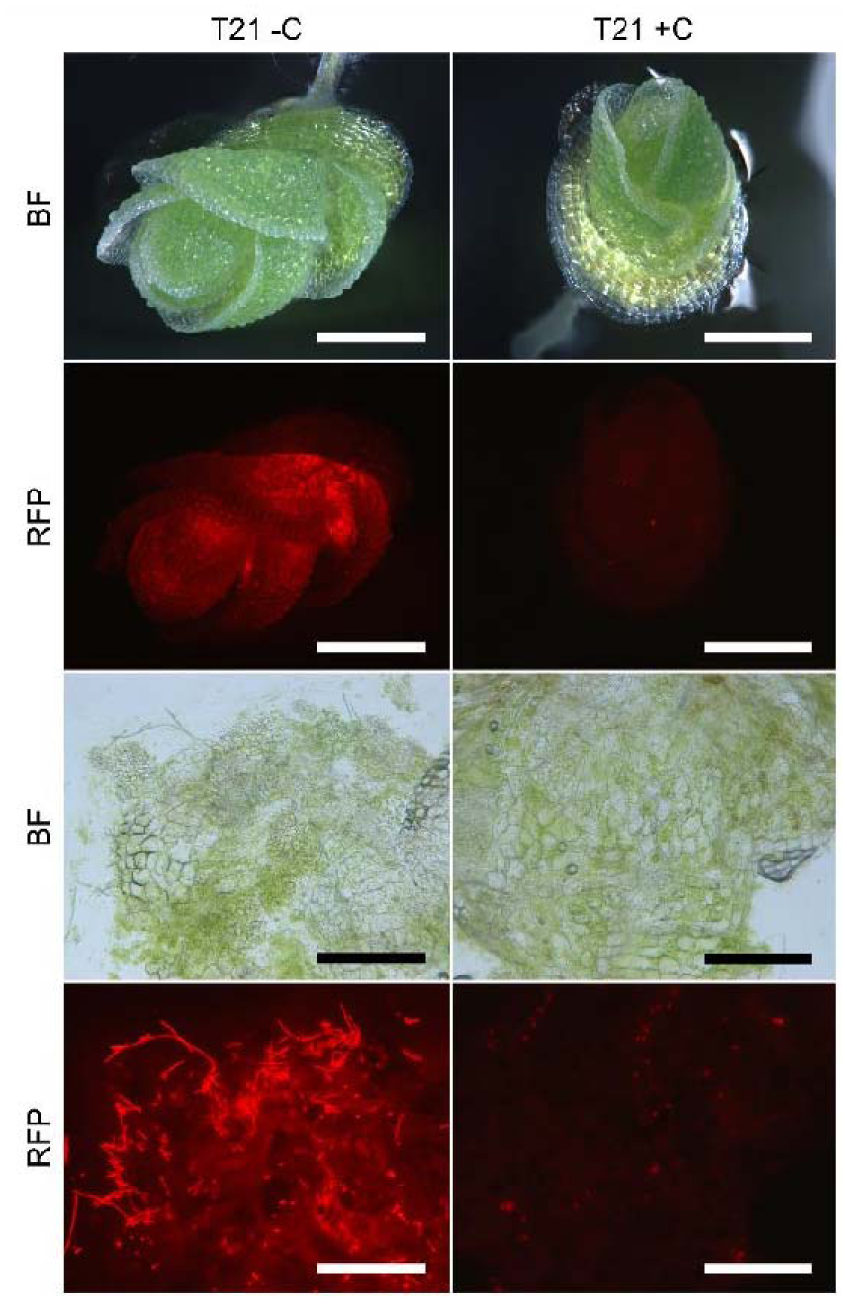
Effect of cornicinine on *A. filiculoides* sporeling development. Top: *A. filiculoides* sporelings 21 days after germination with(out) 500 nM cornicinine (-/+C). Bottom: same sporelings crushed to expose *N. azollae*. BF: bright-field; RFP: red fluorescent protein settings. Scale bars on top and bottom, respectively, correspond to 400 μm and 200 μm.

**Figure S10.**
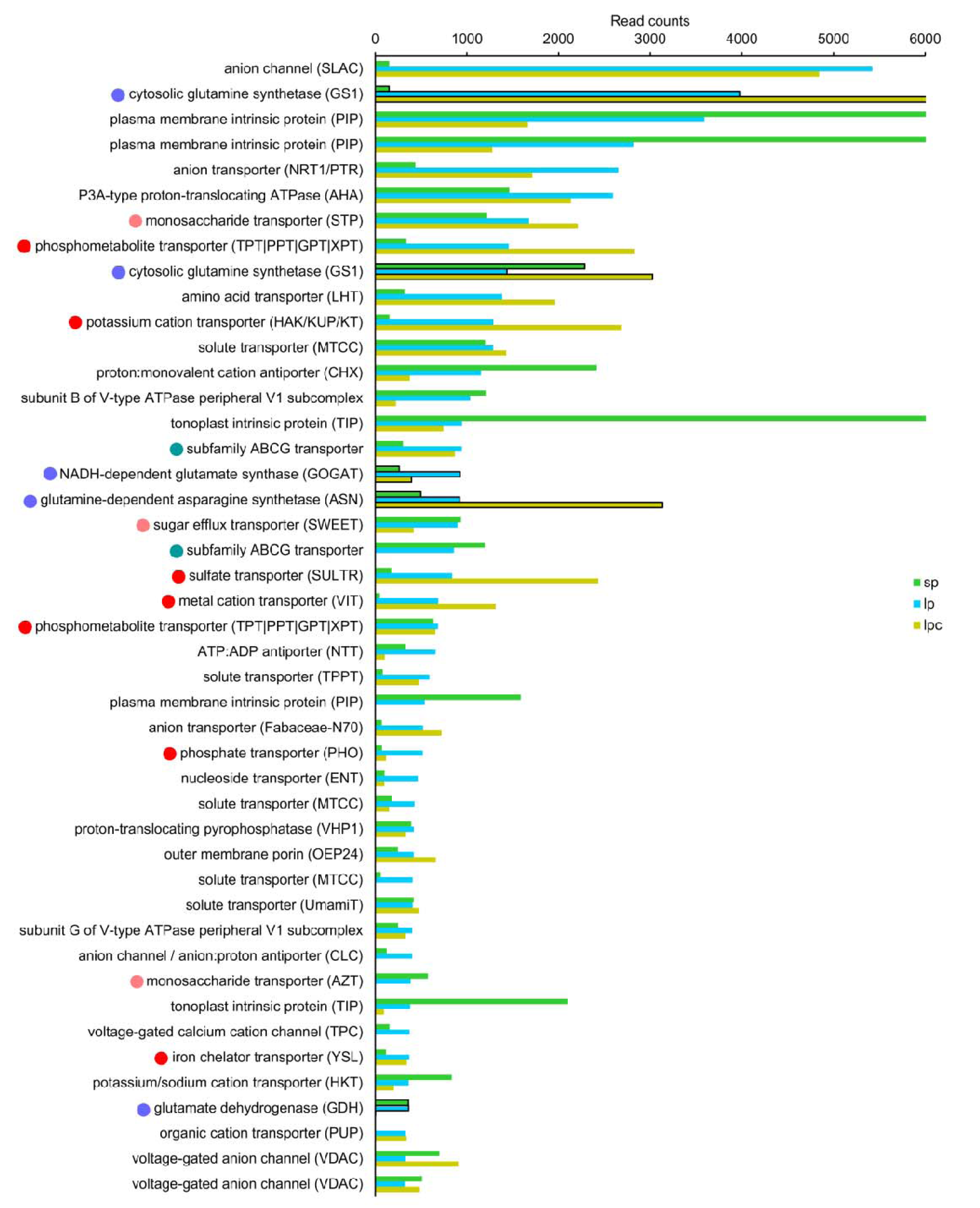
PolyA-tailed transcripts encoding transporters that accumulate most highly in the leaf cavities. Transcripts were ranked according to read counts in the leaf-cavity samples, then selected from the category “transporter” assigned by Mercator annotation. For comparison, the enzymes of ammonium assimilation were also included in the graphic. Averages are shown with n=3, except for lpc where n=2. Samples were: sp, sporophyte; lp, leaf cavities; lpc, leaf cavities from ferns grown on cornicinine. Purple dots highlight enzymes for ammonium assimilation; pink dots highlight sugar transporters; red dots highlight nutrient transporters and; green dots candidates for transport of secondary metabolites.

**Figure S11.**
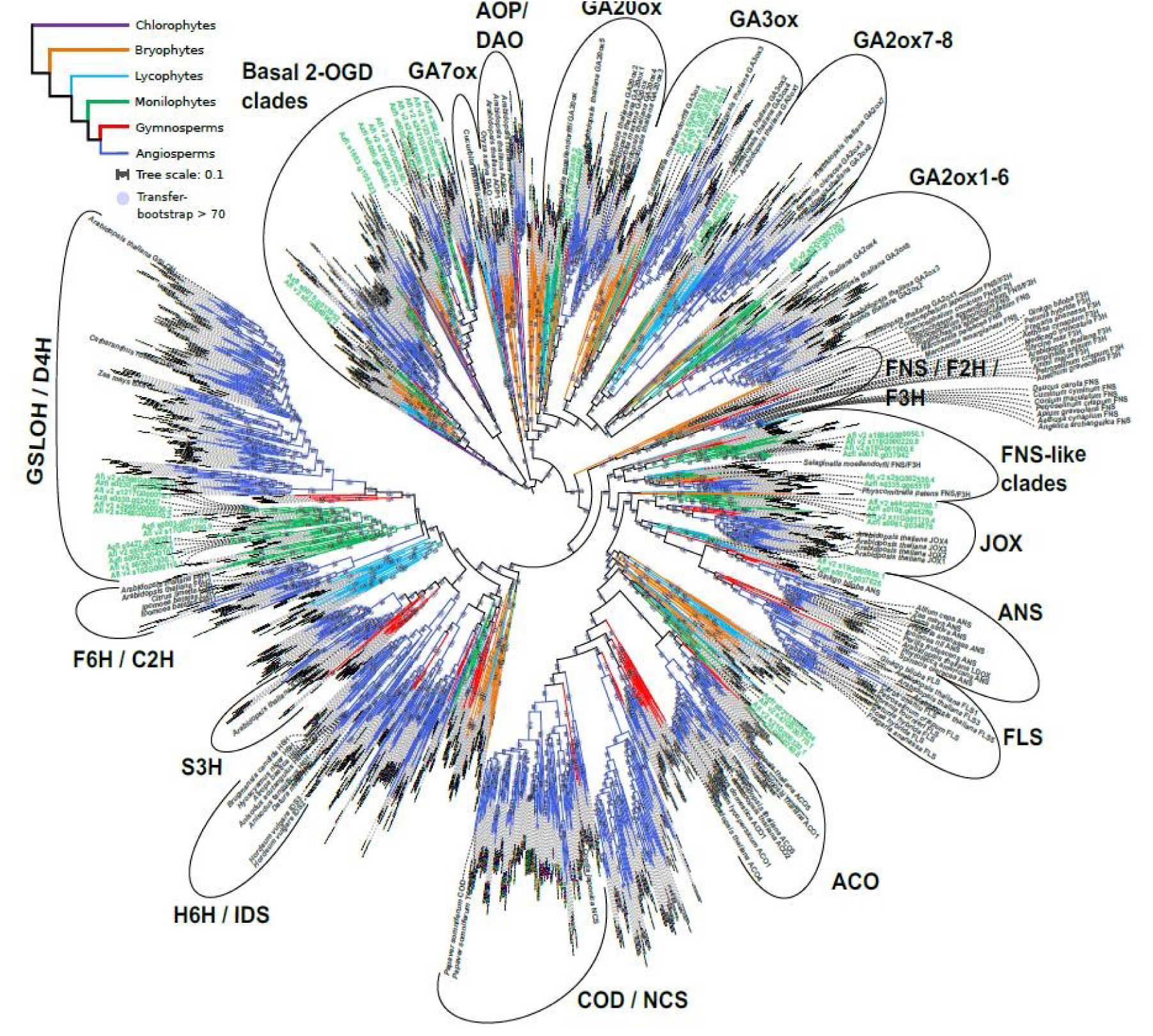
Phylogeny of 2OGD enzymes across land plant lineages. Phylogenies were created of these sequences in the context of class C of 2-oxoglutaratedependent dioxygenases (DOXC, defined in Kawai et al., 2014) using well characterized DOXC enzyme sequences to extract the corresponding orthogroup from the 1kp orthogroup database (Ka-Shu Wong et al., 2019). The orthogroup was sub-sampled and sequences were aligned with MAFFT-einsi (Katoh et al., 2019), and then trimmed using trimAL (CapellaGutierrez et al., 2009). The phylogeny was computed with IQ-tree (Nguyen et al., 2015) with 200 bootstraps. Bootstrap support was calculated as transfer bootstraps (Lemoine et al., 2018). The tree was annotated in iTOL (Letunic and Bork, 2019) and Inkscape.

**Figure S12.**
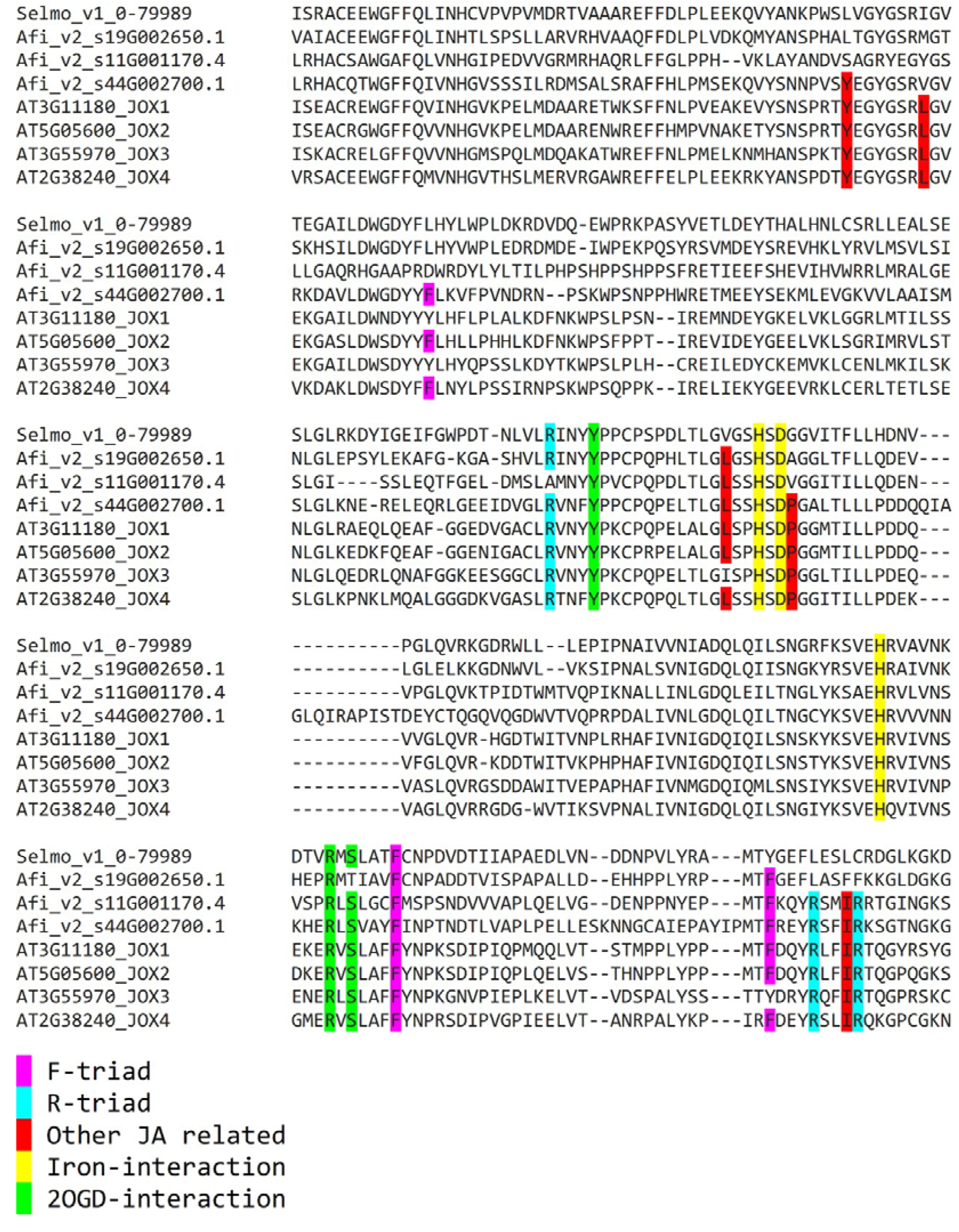
Sequence alignment of the characterized JA-oxidases from Arabidopsis (JOX) and candidate enzymes from *A. filiculoides*, and an enzyme from *Selaginella moellendorfii*. Enzymes are referred to by the gene locus that encodes them. Amino acids highlighted in purple and green constitute the F-and D-triads, respectively, whilst those in red are also known to interact with the JA-substrate. Amino acids highlighted in yellow interact with iron, those in green interact with the substrate 2-oxoglutarate.

## Supporting Files

**Supplemental File 1.** Tables of differentially accumulating transcripts in leaf-pocket preparations with(out) cornicinine, and their expression in sporocarps versus sporophyte. Leaf-pocket profiles with their respective sporophyte control were obtained from sequencing libraries generated using poly-A enriched RNA; read-count normalization was based on read counts of 1100 genes most expressed in the assay (see Materials and Methods). Profiles from the sporocarps versus sporophyte were obtained from sequencing libraries generated from rRNA depleted RNA (dual RNA sequencing); read-count normalization per feature was based on total read counts aligning to the fern genome as in Dijkhuizen et al., 2021, with for each feature the read counts per million reads aligning to the genome. (Supplied as an excel file along with its source data).

